# Low Mutational Load Allows for High Mutation Rate Variation in Gut Commensal Bacteria

**DOI:** 10.1101/568709

**Authors:** Ricardo S. Ramiro, Paulo Durão, Claudia Bank, Isabel Gordo

## Abstract

Bacteria generally live in species-rich communities, such as the gut microbiota. Yet, little is known about bacterial evolution in natural ecosystems. Here, we followed the long-term evolution of commensal *Escherichia coli* in the mouse gut. We observe the emergence of polymorphism for mutation rate, ranging from wild-type levels to 1000-fold higher. By combining experiments, whole-genome sequencing and *in silico* simulations, we identify the molecular causes and evolutionary conditions that allow these hypermutators to emerge and coexist within a complex microbiota. The hypermutator phenotype is caused by mutations in DNA polymerase III, which increase mutation rate by ~1000-fold (a mutation in the proofreading subunit) and stabilize hypermutator fitness (mutations in the catalytic subunit). The strong mutation rate variation persists for >1000 generations, with coexistence between lineages carrying 4 to >600 mutations. This *in vivo* molecular evolution pattern is consistent with deleterious mutations of ~0.01-0.001% fitness effects, 100 to 1000-fold lower than current *in vitro* estimates. Despite large numbers of deleterious mutations, we identify multiple beneficial mutations that do not reach fixation over long periods of time. This indicates that the dynamics of beneficial mutations are not shaped by constant positive Darwinian selection but by processes leading to negative frequency-dependent or temporally fluctuating selection. Thus, microbial evolution in the gut is likely characterized by partial sweeps of beneficial mutations combined with hitchhiking of very slightly deleterious mutations, which take a long time to be purged but impose a very weak mutational load. These results are consistent with the pattern of genetic polymorphism that is emerging from metagenomics studies of the human gut microbiota, suggesting that we identified key evolutionary processes shaping the genetic composition of this community.

## Introduction

Bacteria typically live in multi-species ecosystems [1]. One of such communities is the human gut microbiota, which is key for host health and can host up to 10^13^ bacteria and hundreds of species. Over the last decade, extensive efforts have gone into characterizing the factors (e.g. diet, antibiotics) that shape the species-level composition of this ecosystem (e.g. [2–4]). More recently, it has become clear that there can be abundant genetic diversity within each species colonizing a given individual [5–8]. Importantly, strain level variation is key for multiple phenotypes ranging from antibiotic resistance and virulence to colonization resistance (e.g. [9–11]). However, little is known about how different evolutionary mechanisms – mutation, recombination, migration, genetic drift and natural selection – shape bacterial genetic diversity within the mammalian gut.

Mutation is the raw material for natural selection. The genomic mutation rate of microbes is remarkably constant (0.001/generation) [12]. This suggests that it is shaped by general evolutionary forces, independent of phylogeny or niche [12,13]. Nevertheless, within-species variation for mutation rates can be observed in natural isolates of commensal and pathogenic bacteria (typically to increase mutation rate, *i.e*. mutator clones; e.g. [8,14–17]). Based on the known DNA repair mechanisms, mutators of low to very high strength (10 to >10000-fold mutation rate increases) could in principle emerge and spread [18,19]. However, from the balance between the capacity to adapt (adaptability) and robustness at maintaining current fitness (adaptedness), theory predicts that mutator clones with a strength of 10 to 200-fold should be the most common (e.g. [20,21]). Indeed, when mutator emergence was observed *in vitro*, very strong effect mutators (>300-fold) have not been detected [22–24]. However, in the experiments above, a bacterial lineage evolved in the absence of the multitude of biotic interactions that occur in natural ecosystems, such as the mammalian gut microbiota. In this ecosystem, biotic interactions can rapidly fluctuate [25,26], either due to changes in community composition or migration of novel lineages, often caused by community perturbations as dietary changes or antibiotic intake [2,3,27]. Such environmental fluctuations are theoretically predicted to lead to mutation rate variation [21,28]. Yet, we have limited knowledge of within-host mutation rate variation [14–17,29] and its temporal dynamics within a diverse multi-species community [8].

Here, we followed the long-term evolution of a commensal *E. coli*, when invading the complex ecosystem of the mouse gut after a perturbation caused by antibiotic treatment. Using an experimental evolution approach, we tracked the evolution of two defined *E. coli* lineages. The real-time evolution in the gut revealed the coexistence of lineages with mutation rates ranging from wild-type levels to 1000-fold higher. We determine both the molecular causes and the evolutionary conditions that allow such strong mutators to emerge and persist within a complex microbiota. At the molecular level, we show that mutations in the proofreading (α) and catalytic (ε) subunits of DNA polymerase III cause a 1000-fold mutation rate increase and improve hypermutator fitness, respectively. At the evolutionary level, we show that the population dynamics of the mutator lineages and their pattern of molecular evolution can be explained if deleterious mutations have very weak effects, on the order of ~0.01-0.001%. Such effects are 100 to 1000-fold lower than the current estimates from mutation accumulation experiments in laboratory environments (1-3%) [30,31] and would be unmeasurable by direct competitive fitness assays. Despite the large number of deleterious mutations observed in the mutators, we can identify multiple beneficial mutations. However, no fixations were detected, neither of specific alleles nor of changes at the functional level (gene/operon). This suggests that negative frequency-dependent or fluctuating selection shape the trajectories of most beneficial mutations (rather than constant positive Darwinian selection). The data reveal that the evolution of commensal bacteria within the mammalian gut is consistent with the nearly neutral theory of molecular evolution combined with partial hitchhiking events caused by the increase in frequency of linked beneficial mutations *(i.e*. genetic drafts) [32,33].

## Results and Discussion

### Emergence and maintenance of mutation rate variation in a gut commensal strain

We used a short-course antibiotic treatment (8 days with streptomycin) to induce a perturbation of the microbiota and break colonization resistance, which allows us to study the evolution of new *E. coli* strains as they invade and colonize the mouse gut. With this setup, we colonized four mice with two commensal strains of *E. coli* (marked with two fluorescent-markers: YFP and CPF; both streptomycin-resistant) and followed their dynamics for 190 days (~3600 generations, assuming 19 generations/day; Fig. 1A) [34,35]. Throughout the entire experiment, both strains coexisted in all mice and *E. coli* generally maintained a population size of >10^7^ CFU/faeces gram (Fig. S1), *i.e*. the typical size of its ecological niche in this host [36].

**Figure 1.**
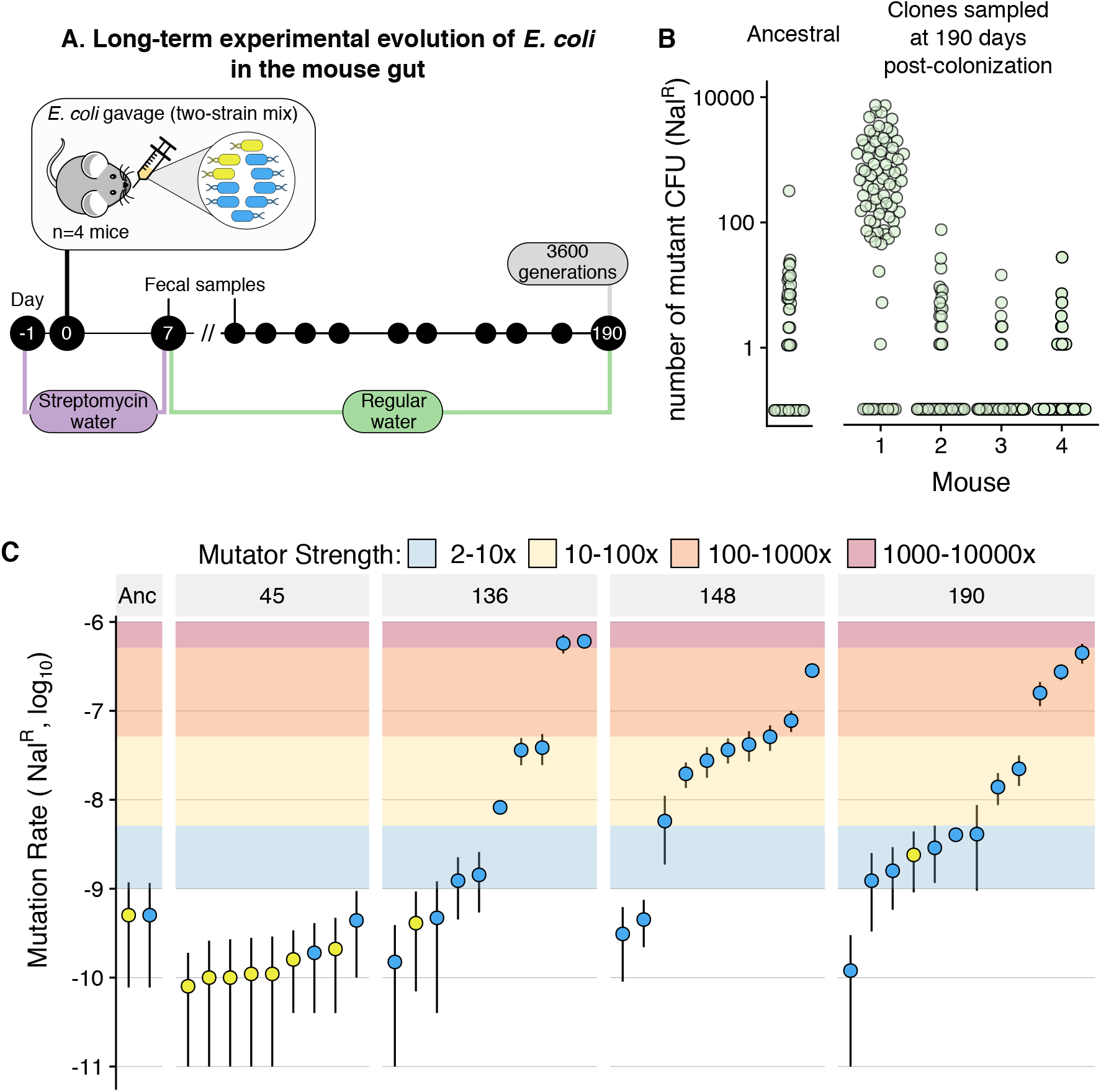
Multiple clones with variable mutation rates (up to 1000-fold higher than ancestral) emerge and coexist during long-term adaptation to the mouse gut. **A)** Scheme of the experimental design used to follow the long-term adaptation of *E. coli* to the mouse gut. **B)** Screen for mutation rate variation shows that mutators emerged in mouse 1. 83-90 clones, isolated from each mouse at day 190 post-colonization, were grown overnight and plated in nalidixic acid. The number of resistant colonies obtained for each clone is shown. The same procedure was carried for 96 replicates of the ancestral clones. **C)** Large scale variation for mutation rate emerges in mouse 1. Mutation rates (towards nalidixic acid resistance), measured through fluctuation tests, for multiple clones randomly isolated at different time points from mouse 1. Each point corresponds to one clone. Error bars represent 95% confidence intervals of 10 independent growths (non-overlapping bars indicate significant differences). Blue and yellow points correspond to clones from the CFP and YFP backgrounds. Coloured areas show the corresponding mutator strength (i.e. fold increase in mutation rate, relative to the ancestral).

As mutation rate is a key evolutionary parameter that can evolve, both *in vitro* and *in vivo*, we screened a sample of clones from each mouse (n~90 per mouse) for potential changes in mutation rate (Fig. 1B). In one of the four mice, we detected clones with increased mutation rate. To understand the magnitude and dynamics of the mutation rate increase in this host, we performed Luria-Delbrück fluctuation assays for several clones sampled at days 45, 136, 148 and 190 and measured their mutation rate towards resistance to nalidixic acid (n=10 independent growths per clone; Fig. 1C). At day 45 none of the clones exhibited a mutation rate significantly different from the ancestral (5×10^−10^; 95% confidence intervals for all clones overlap with those of the ancestral). However, from day 136 onwards, mutation rate varied by three orders of magnitude, ranging between 4.7×10^−10^ and 6×10^−7^. Similar levels of mutation rate variation were observed at days 148 (range: 3.1×10^−10^ to 2.8×10^−7^) and 190 (1.2 x10^−10^ to 4.5×10^−7^). Thus, non-mutator clones coexisted with both strong hypermutators (mutation rates up to 1200-fold higher than the ancestral) and with more commonly observed hypermutators (10-100-fold increases in mutation rate). This indicates that multiple mutators had emerged either simultaneously or sequentially. To the best of our knowledge, these results represent the first observation of spontaneous emergence of 1000-fold mutator clones and of long-term maintenance of large scale mutation rate polymorphism (>1000 generations, *i.e*. >54 days).

These observations raised two main questions: What is the genetic basis for the observed mutation rate increase? What are the evolutionary conditions that could allow for the emergence of 1000-fold mutators and for the long-term maintenance of mutation rate polymorphism?

### Mutations in the proofreading and catalytic subunits of DNA polymerase III cause hypermutability and improve mutator fitness, respectively

To determine the cause of the observed mutation rate variation, we carried out whole-genome sequencing of 18 clones, 13 mutators and 5 non-mutators, isolated between day 136 and 190 post-colonization. All mutator clones shared 45 mutations, none of which was present in the isolates that maintained the ancestral mutation rate (Table S2). Among these, we identified a single non-synonymous mutation in the gene *dnaQ*, which could increase mutation rate by 1000-fold. *dnaQ* encodes the proofreading (ε) subunit of DNA polymerase III [37], the main DNA polymerase in *E. coli*. Following their divergence, mutators accumulated mutations in several other genes that may affect mutation rate (Fig. S2; none of the genes was mutated in the non-mutators; Table S4). Strikingly, these included several mutations in two other subunits of DNA polymerase III: the catalytic (α) subunit (encoded by *dnaE;* mutated in 11 clones) [38] and in the θ subunit *(holE;* two clones) [39]. The α, ε and θ subunits physically interact to form the core of DNA polymerase III (Fig. 2A) [19,40], and non-synonymous mutations in *dnaQ* were previously shown to cause mutation rate increases ranging to >5000-fold [18]. Mutations in *dnaE* and to a smaller extent in *holE* were also shown to have anti-mutator effects when linked to *dnaQ* mutations [41,42]. This led us to hypothesize that a first mutation in *dnaQ* raised the mutation rate, which was reduced by subsequent mutations in *dnaE*, thus leading to the observed mutation rate polymorphism. To query if the *dnaQ* mutation caused the hypermutator phenotype and if *dnaE* could act as a modifier, we engineered single and double mutants with the *dnaQ* mutation (L145P; C<u>T</u>C → C<u>C</u>C) and a mutation in *dnaE* (T771S; <u>A</u>CG → <u>T</u>CG) and measured their mutation rate. We generated ~20 clones carrying the DnaQ^L145P^ allele, as *dnaQ* hypermutators can exhibit high mutation rate variation, possibly due to the rapid emergence of suppressors [42,43]. The *dnaQ* clones indeed showed a very high mutation rate, reaching 3000-fold increases for both nalidixic acid and rifampicin (Fig. 2B). The distribution of mutation rate estimates is bimodal, with a group of clones showing ~200-fold increase in mutation rate and another showing an increase of ~1000-fold (none of the 95% CIs overlap with the ancestral; Table S1). Given that we observe mutation rate increases of ~1000x for clones isolated from mice and due to the known possibility of suppressors emerging during the engineering process [42], we suggest that the actual effect of the *dnaQ* mutation is to increase mutation rate by ~1000 fold.

**Figure 2.**
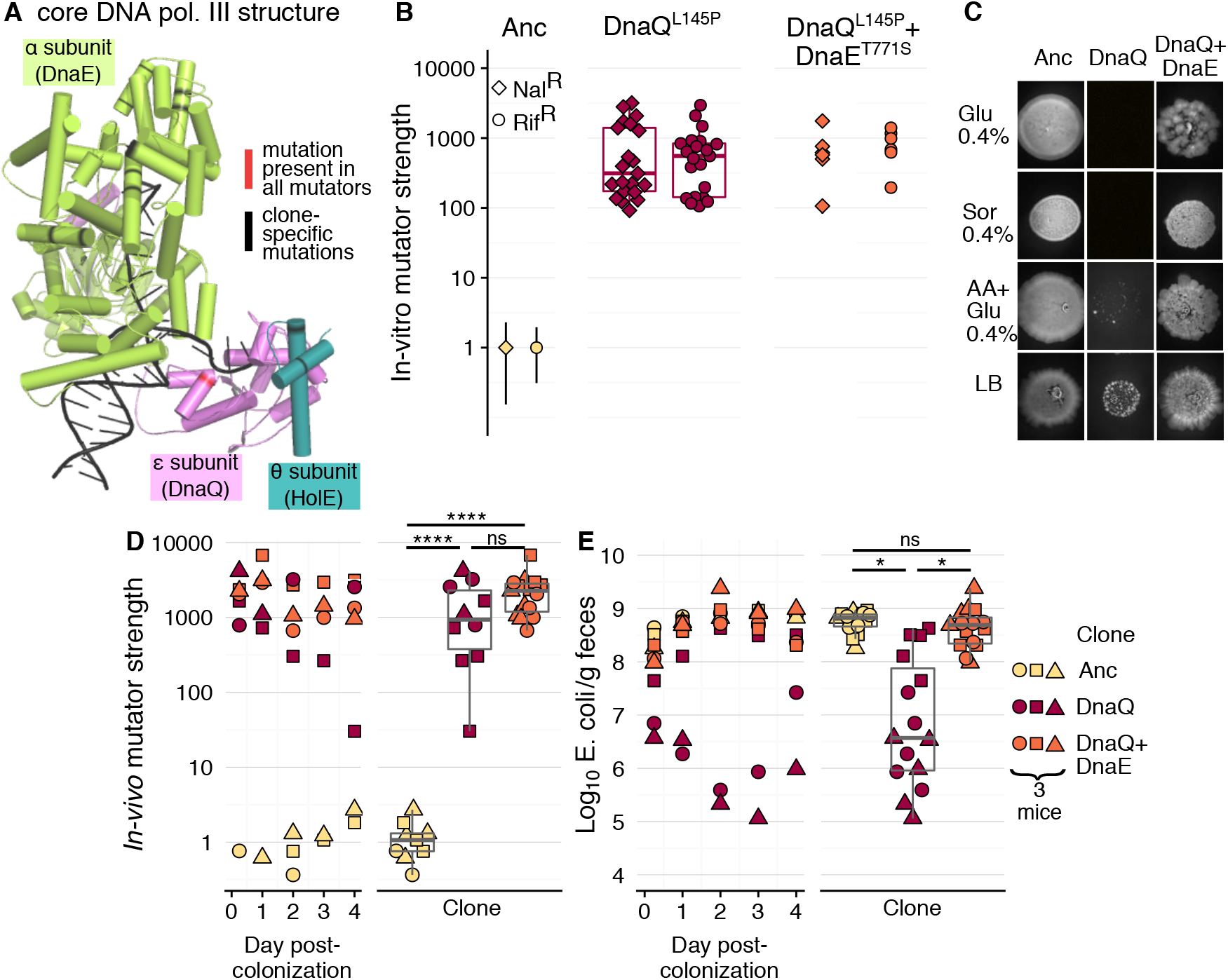
Mutations in DNA polymerase III both drive mutation rate increase by 1000-fold (DnaQ^L145P^; ε subunit) and rescue fitness of hypermutator clones (DnaE^T771S^; α subunit). **A)** Structure of the core DNA polymerase III (PDB ID: 5M1S), with each subunit in a different colour and non-synonymous mutations highlighted in red or black. **B)** *In vitro* fluctuation tests were carried for the ancestral clone, the single mutant DnaQ^L145P^ and the double mutant DnaQ^L145P^+DnaE^T771S^ and the double mutant (21 and 5 independent clones were tested for each mutant, respectively). Each point shows the mutation rate estimate relative to the ancestral (*in vitro* mutator strength) for an independent clone (10 replicates per clone for ancestral and double mutant; 5 replicates per clone for DnaQ^L145P^). For ease of visualization, 95% confidence intervals are only shown for the ancestral, but none of the 95% CI of either mutant is overlapping with those of the ancestral (See Table S1 for mutation rates and 95% CI). **C)** *in vitro* phenotypic growth capacity of the ancestral clone, the single mutant DnaQ^L145P^ and the double mutant DnaQ^L145P^+DnaE^T771S^ (spot assay; all inocula had the same OD_600nm_). **D-E)** *In vivo* dynamics and summary boxplots of the mutation frequency relative to the ancestral (*in vivo* mutator strength; D) and *E. coli* CFUs per gram of faeces (E).

In contrast to our hypothesis, the DnaE^T771S^ mutation did not cause any significant change in the mutation rate, neither on the ancestral nor on the DnaQ^L145P^ backgrounds (Fig. 2B and Fig. S3). This suggests that the *dnaE* mutation did not act to decrease mutation rate. However, the independent occurrence of different alleles in the α subunit of DNA polymerase III (Fig. 2A and Fig. S2) strongly suggests that mutations in *dnaE* could be beneficial. Consistent with this hypothesis, we observed that DnaE^T771S^ strongly improves growth of the DnaQ^L145P^ mutant (Fig. 2C and S4). Remarkably, the DnaQ^L145P^ mutation is lethal in minimal media with a single carbon source (either glucose or sorbitol). Its null fitness can be rescued by complementing the minimal media with amino acids or in LB, which allow the DnaQ^L145P^ mutant clones to achieve slow but visible growth (Fig. 2C). However, the effect of the *dnaE* mutation is stronger than that of the environment, as it allows growth of the double mutants to recover to levels similar to the ancestral (non-mutator) clone, across all media.

As mutation rate and spectrum can depend on the environment (e.g. [44]), we sought to determine the mutation rate of the *dnaQ* and *dnaQ*+*dnaE* mutant strains *in vivo*. The classical Luria-Delbruck *in vitro* assay, designed to avoid selection, is inappropriate for estimating the mutation rate in the gut. However, we can use Haldane’s mutation-selection balance principle to estimate the *in vivo* mutation rate of these clones. For that, we colonized mice with the ancestral, the *dnaQ* mutant and a double *dnaQ+dnaE* mutant and measured the equilibrium frequency of rifampicin-resistant (Rif^R^) clones (n=3 mice for each genetic background). Under mutation-selection balance, the frequency of resistant mutants is directly proportional to the mutation rate [45]. Thus, the ratio of the Rif^R^ mutation frequency in the mutators, relative to the ancestral, provides a direct estimate of mutator strength. The mutation frequency of either the single *dnaQ* mutant or the double *dnaQ+dnaE* mutant is ~1000-fold higher than observed for the ancestral, but not significantly different between mutants (Fig. 2D, see also Fig. S5 for comparison between mutation frequency *in vivo* and in an *in vitro* propagation; linear mixed model with Tukey test: clone effect: **χ**^2^_2_=31.89, *p*<0.0001; ancestral vs *dnaQ*: *t*=−12.81, *df*=6, *p*<0.0001; *ancestral* vs *dnaQ+dnaE*: *t*=−14.94, *d*f=6, *p*<0.0001; *dnaQ* vs *dnaQ+dnaE*: *t*=−1.47, *df*=6, *p*=0.37). When comparing the niche size that each strain occupies in the mouse gut, we find that the abundance of the *dnaQ* mutant (10^5^-10^6^/feces gram) is significantly lower than that of the ancestral (10^8^/feces gram; linear mixed model with Tukey test: clone effect: **χ**^2^_2_=10.16, *p*=0.006; ancestral vs *dnaQ*: *t*=−3.74, *df*=6, *p*<=0.022). Remarkably, the fitness defect of *dnaQ* is fully rescued by the *dnaE* mutation (Fig. 2E), as the bacterial loads for the double mutant are similar to those for the ancestral (>10^8^/feces gram; *ancestral* vs *dnaQ+dnaE*: *t*=0.23, *df*=6, *p*=0.97; *dnaQ* vs *dnaQ+dnaE*: *t*=−3.51, *df*=6, *p*=0.029). These results are similar to those observed *in vitro*, when the strains grow in different nutrient sources. Taken together, our results establish that the *dnaQ* mutation massively increases mutation rate both *in vitro* and *in vivo*, albeit at a fitness cost. This cost can be recovered by a beneficial mutation in *dnaE*, without strong effects on mutation rate. The causes of mutation rate polymorphism are likely to be complex, as most mutator clones are mutated in ≥3 genes that can affect mutation rates (e.g. a single clone can have mutations in *dnaQ*, *dnaE, recF, katE* and *sodB*; see Fig. S2).

### Survival of hypermutators indicates that deleterious mutations are of small effect

The emergence and long-term maintenance of mutators with a 500-1000x increase in mutation rate raises a conundrum: How can such strong mutators avoid a catastrophic fitness decline caused by Muller’s ratchet [46,47]? Clonal populations with a genomic mutation rate reaching 1 new mutation per generation should quickly accumulate many deleterious mutations and should show severe impediments on fixing adaptive mutations (given current estimates for the fitness effect of deleterious mutations of 1-3%; [30,31,48]). Due to the accumulating mutational load, such strong mutators should continuously decrease in fitness (*i.e*. Muller’s ratchet), in a potentially inescapable path towards extinction [49,50]. To understand under which conditions mutators could avoid this fate, we simulated Muller’s ratchet (see methods) under many combinations of the deleterious mutation rate *(U_d_)* and the negative effects of mutations *(s_d_)*. We started with the simplest model of the ratchet, ignoring compensatory or beneficial mutations. We assumed a population of size 10^6^ (similar to *E. coli*’s population size in the gut; Fig. S1) and quantified fitness and the number of accumulated mutations after 1000 and 2000 generations (~55 and 110 days, similar to the time during which mutator maintenance was observed).

We first consider *s_d_* values of 1%, similar to those typically estimated from *in vitro* mutation accumulation experiments with *E. coli* [30,31]. Under this *sd* value, the fitness of lineages with very high mutation rates (*U_d_*>0.3) declines to <0.5 within 1000 generations. Such lineages should therefore be outcompeted by clones with a wild-type mutation rate (whose fitness remains unaltered, independently of *s_d_*; Fig. 3A and Fig. S6–7). Remarkably, when *s_d_*≤0.01%, 100-fold smaller than previously assumed, the simulations predict that: 1) hundreds of *de novo* deleterious mutations should accumulate; 2) strong mutators can survive for periods of ~100 days. These general predictions are robust to the presence of beneficial mutations (see below and [51]). To determine if these expectations are met in the evolving lineages, we analyzed the genomes of the 13 mutator and 5 non-mutator clones. The mutator and non-mutator clones followed independent evolutionary paths, with non-mutators acquiring only 4 to 5 mutations, while mutator clones accumulated between 164 and 658 mutations (Fig. 3B). Such a large variation in the number of mutations is hard to explain, as shown in simulations where a lineage with a single mutation rate is segregating (Fig. 3A and S6–7). Furthermore, with *sd* values between 0.1% and 1%, lineages with ~600 mutations can only be detected after 2000 generations (when Ud>0.5), and their fitness declines to between <0.01 and 0.3 (Fig. S6–7). Such strong fitness declines would require equally strong beneficial mutations for hypermutator maintenance. However, the fitness effects of strongly beneficial mutations in the mouse gut have been shown to vary between 2 and 12% [35,52], which would be insufficient to maintain lineages with 600 mutations. Moreover, if strong enough beneficial mutations were available, these would fix very rapidly [53] which is at odds with the observed frequency of beneficial mutations (see below). In contrast, if the effects of deleterious mutations are at least two orders of magnitude lower, s¿<0.01%, lineages with ~600 mutations are already predicted at 1000 generations and their fitness remains >0.94, being very close to the fitness of lineages with 200 mutations (always >0.98). Similar results are obtained in simulations with a continuous distribution of fitness effects, where *s_d_* is exponentially distributed (Fig. S8), rather than constant. As expected from these results, in simulations in which clones with distinct mutation rates compete, mutation rate polymorphism can only be maintained if *s_d_*≤0.01%. This result is largely unaffected when including beneficial mutations (Fig. S9).

**Figure 3.**
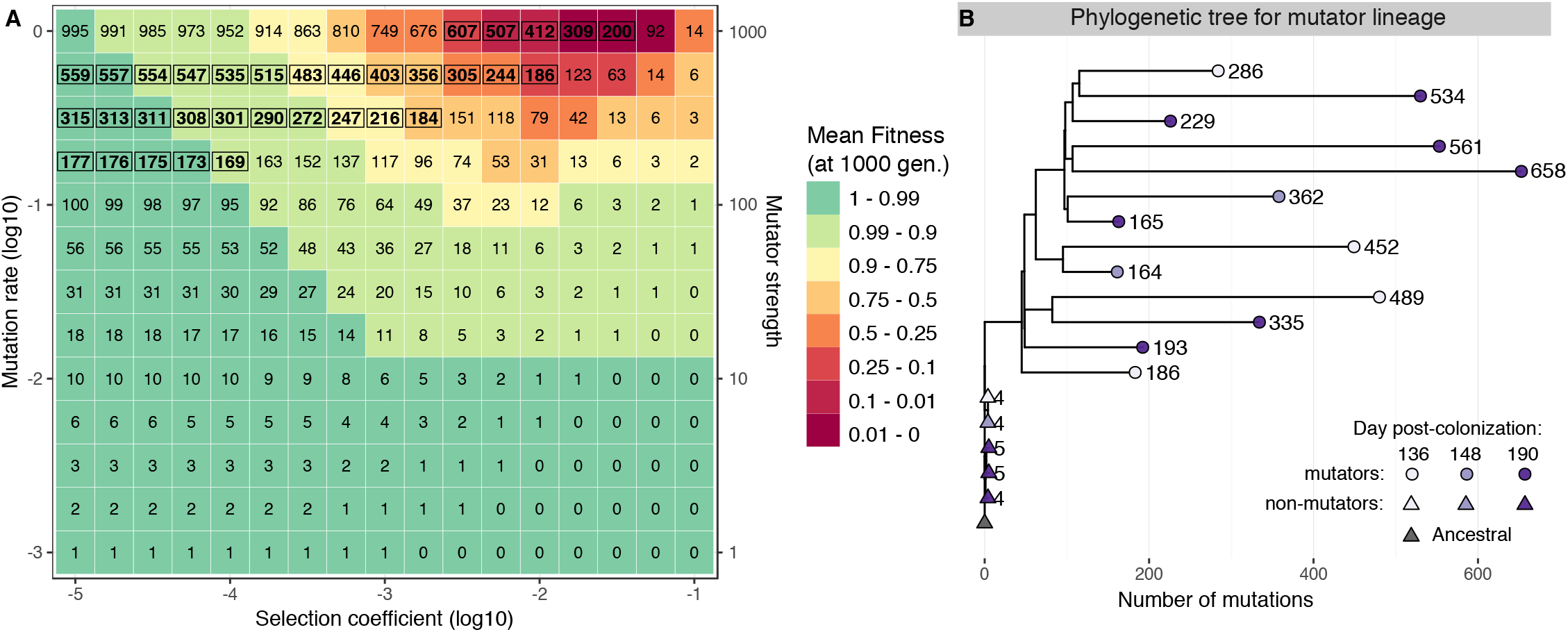
Strong, intermediate and weak mutators can escape Muller’s ratchet if the effects of deleterious mutations are much weaker (<0.01) than current estimates (1%). **A)** Mean fitness and mean number of mutations in Muller’s ratchet simulations with deleterious mutations of fixed effect and a clonal population size of 10^6^ (10 simulations). Colour gradient indicates the mean fitness after 1000 generations, which starts at 1 and declines to different levels, depending on the combination of selection coefficient (*s_d_*) and mutation rate (*U_d_*). Numbers in bold and inside a box indicate the parameter space for which there is a fit between the observed and the simulated number of mutations (see methods). The observed variation in the number of mutations is only possible if *s_d_*<10^−4^ and if there is mutation rate variation. **B)** Mutation accumulation in wild-type and mutator clones shows extremely large variation in the rate of molecular evolution for new lineages emerging within the gut microbiota. Phylogenetic tree for 18 clones from mouse 1 and the ancestral, generated from whole-genome sequence data. Branch lengths and tip labels indicate the number of accumulated mutations. Tip point colours indicate the day post-colonization at which different clones were isolated.

Thus, the theoretical expectations constrained by the experimental observations (*in vivo* mutation accumulation and long-term maintenance of mutation rate polymorphism; Fig. 1 and 3), are consistent with *s_d_* values much lower than what has been inferred from classical *in vitro* mutation accumulation experiments (*s_d_*~1 to 3%), including the recent estimates from single cells (*s_d_*=0.3%) [54]. Such weak effects of deleterious mutations, inferred here to explain the dynamics of the mutators in a natural environment, are much more consistent with typical estimates of the fitness effect of deleterious mutations obtained from analysis of natural polymorphism (e.g. for *Drosophila, s_d_*~10^−5^ − 10^−4^ when estimated from natural polymorphism, 100-1000 fold lower than inferred from mutation accumulation data; [55,56]). Moreover, despite the gap in knowledge of the causes of polymorphism in commensal bacteria [57], its high level indicates that deleterious mutations should typically have small effects [58,59]. Remarkably, by analysing polymorphism patterns in species of the human gut microbiome (including *E. coli*), Garud et al. [5] recently estimated *s_d_*/*μ* = 10^5^ (*μ* - mutation rate per site). This implies a *s_d_* ~ 10^−5^, which is fully consistent with our results (with *μ* = 10^−10^, as estimated for *E. coli* from mutation accumulation experiments coupled with whole-genome sequencing; [60]).

### Partial selective sweeps structure the high polymorphism of a commensal strain evolving in the mouse gut

Whereas weak effect deleterious mutations can explain the observed molecular evolution patterns, they cannot explain mutator invasion. In *in vitro* evolution experiments, mutators have been seen to invade through hitchhiking with beneficial mutations [23,61,62]. Given the fitness defect of the *dnaQ* mutation described above (*in vitro* and *in vivo*) and that beneficial mutations can emerge rapidly during adaptation to the mouse gut [34,35,52,63], we propose that hitchhiking is also the most likely route driving the observed increase in mutation rate. Thus, we sought to find signatures of adaptation in the mutator clones. First, we computed the ratio of non-synonymous to synonymous changes, dN/dS, to detect whether there was evidence of either purifying (dN/dS<1) or positive selection (dN/dS>1) [64]. Consistent with the above modelling inference, where a large number of very slightly deleterious mutations accumulated in the mutators, dN/dS showed a weak signal of purifying selection. (dN/dS < 1 in 3/13 clones; P<0.05; Fig. S10). Furthermore, no sign of positive selection could be found using this statistic (a well-known difficulty for clonal populations).

However, due to the power of our experimental design, where the same colonizing lineages evolve independently in genetically identical animals under the same diet, one can seek to identify beneficial mutations via mutational parallelism. For this, we isolated 3-4 clones per mouse (from the remaining 3 mice) at day 190 post-colonization and carried out whole-genome sequencing. We then identified genes or operons that had been repeatedly mutated across hosts (i.e. parallelism; Fig. S11) as likely targets of natural selection. Using this approach, we identified 12 targets, 8 of which were also mutated in the mutator clones. Of these, *ptsP, frlR* and *dgoR* were mutated in all the mutators (*i.e*. in the mutator common ancestor), while 5 other targets were mutated in at least one of the mutator clones (Fig. 4).

**Figure 4.**
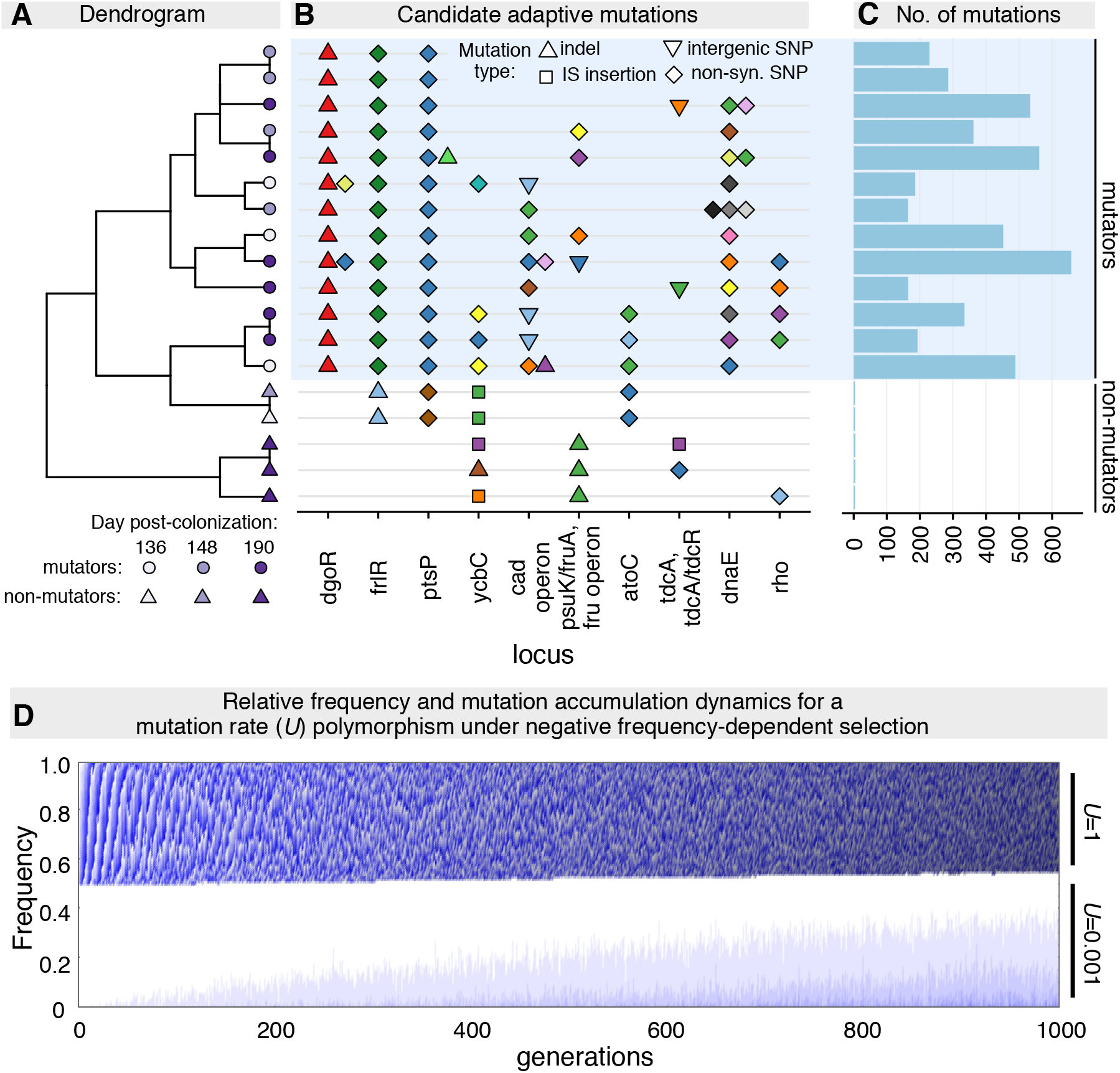
Multiple adaptive mutations accumulate in mutator and non-mutator clones without fixing, highlighting the importance of negative frequency-dependent selection. **A)** Clustering of clones depending on which candidate targets of adaptation are mutated. **B)** Candidate mutational targets of adaptation that are mutated in each clone (different colours indicate different alleles). All targets that showed parallelism across mice are included plus *dnaE* and *rho*, which were identified from parallelism across mutator clones and for which we have independent evidence of adaptation. 39 additional targets were identified through parallelism across mutator clones (Table S3). **C)** Number of mutations accumulated per clone. **D)** Negative frequency-dependent selection can maintain stable coexistence between two clones with largely different molecular evolution patterns (mutation rate, *U*, differing by 1000-fold). Within 1000 generations, the clone with *U*=1 accumulates up to 1000 mutations, whereas the clone with *U*=0.001 accumulates only up to 3 mutations. Darker tones of blue and grey indicate larger numbers of mutations. Simulation parameters: N=10^6^; *s_d_*=10^−5^; strength of negative frequency-dependent selection: 0.1.

Most parallel targets in the mutators have a metabolic function and some have been previously observed in other mouse gut colonization experiments. Specifically, while *ptsP* is a novel target of adaptation, *frlR* and *dgoR* have been previously identified as targets of adaptation to the gut of streptomycin-treated immunocompromised or wild-type hosts, respectively [34,63]. *frlR* and *dgoR* are negative regulators of the *frl* and *dgo* operons [63,65,66], which are involved in carbon metabolism, namely of fructoselysine and galactonate (respectively). *ptsP* encodes a phosphoenolpyruvate-protein phosphotransferase, which is thought to be involved in the response to nitrogen availability and to several stresses (e.g. salt, pH) [67–69], both important pressures in the gut [70–72]. Of these three, *dgoR* is the most likely candidate for the adaptive mutation with which the *dnaQ* mutator allele could have hitchhiked, since *dgoR* is mutated in clones from all other mice, but only in the mutator clones from mouse 1. Furthermore, *dgoR* is the main target of adaptation when a uropathogenic *E. coli* colonizes laboratory mice of a different genetic background [63]. This strongly suggests that mutations in *dgoR* are strongly adaptive and such adaptation transcends the specific *E. coli*-mouse strain combinations. Interestingly, the mutations targeting *dgoR* and *frlR* (in this and in previous studies) often involve IS insertions, indels or nonsense mutations. These are likely to lead to loss of function of the negative regulators and thus, overexpression of the regulated operons. Such adaptive mutations are expected if galactonate and fructoselysine are present in the gut as limiting resources for *E. coli*. This form of adaptation has been previously shown to occur for the case of galactitol [73].

After divergence from their common ancestor, the mutators acquired potentially adaptive mutations in the *cad* operon, *ycbC, atoC, psuK/fruA* and *fruB*, and *tdcA/tdcR*. Of these targets, mutations in the last two have previously been observed during adaptation of *E. coli* to the gut of immune-competent or immunocompromised mice [34,36]. Many of the adaptive targets are functionally important for regulating carbon (*dgoR, frlR, atoC* and, potentially, the *cad* operon) [67,74–76] or nitrogen metabolism (*ptsP* and *tdcA/tdcR*) [69,77,78], stress resistance (*ptsP, cad* operon, *ycbC* and, potentially, *atoC*) and peptidoglycan biogenesis (*ycbC*) [67,74–76]. Interestingly, *ptsP, atoC* and the *cad* operon appear to be involved in regulating both nutritional competence and stress resistance, which suggests that such mutations may modulate a potential trade-off between nutritional competence and stress resistance, which is well described in *in vitro* experiments and which is also thought to be important *in vivo* [71,72,79]. Together with *dgoR*, the *cad* operon is the only other mutational target that occurs in all other mice, but is specific of mutator clones in mouse 1.

As the mutator clones have largely independent evolutionary histories (long terminal branches in Fig. 2B), the concept of parallel evolution can also be used to identify adaptive targets that were specific to the mutator lineages. However, as the mutation rate is extremely high, parallel mutations can occur just by chance. Thus, we used simulations and conservative filters to identify genes for which the observed number of non-synonymous SNPs is significantly higher than what would be expected if the observed mutations were randomly distributed (see methods; [80]). 41 genes could be detected with this method, the majority of which are involved in nitrogen or carbon metabolism (Table S3), which is in line with the above results. Among these, we found *dnaE* and *cadC*, which had been identified as adaptive via direct experiments or through parallelism across hosts, suggesting that the method is capable of identifying beneficial mutations. Additionally, one of the mutational targets is *rho*, which has also been detected during adaptation of adherent-invasive *E. coli* to the mouse gut [81] and appears in a non-mutator clone.

Among the tens of adaptive mutations that we identified, none reached fixation. Indeed, even after 190 days of evolution (~3600 generations), no complete sweep could be observed, neither of a single mutation (hard sweep) nor of multiple, functionally equivalent, lineages (soft sweep). This indicates that most beneficial mutations in this ecosystem are only conditionally beneficial. This suggests that a more complex form of selection, such as negative frequency-dependent or fluctuating selection, can maintain the observed molecular evolution polymorphism. A dominant role for this form of selection is in complete agreement with three observations: lack of fixation of adaptive mutations (Fig. 4 and Fig. S11), long-term maintenance of polymorphism for mutation rate (Fig. 1) and at a neutral locus (YFP/CFP; Fig. S1). As we have previously observed negative frequency-dependent selection acting on beneficial mutations in the mouse gut [35], we simulated a population with two clones, differing in mutation rate by 1000-fold (*U*=1 and *U*=0.001), whose genotypes include a locus that is under negative frequency-dependent selection of strength 0.1. These clones also accumulate recurrent deleterious mutations of effect *s_d_*=0.001%. Under these conditions, the two clones can coexist for at least 1000 generations at a stable relative frequency of ~50% (Fig. 4D) and with markedly different patterns of molecular evolution. Similar to what we observe experimentally (Fig. 4C), negative frequency-dependent selection leads to stable coexistence of individuals carrying up to 1000 mutations with those carrying just 3 or 4. Importantly, such stable coexistence would not be expected if adaptive mutations were unconditionally beneficial (Fig. S9).

## Conclusions

The emergence and temporal dynamics of mutator populations can provide information on key evolutionary parameters where direct experimental measures may be difficult to obtain. Of key relevance, these can allow the fitness effects of deleterious mutations to be inferred [30,31]. Here, we studied the emergence of multiple mutator lineages during colonization of the mouse gut, including a hypermutator with up to 1000-fold increase in mutation rate. The pattern of molecular evolution is consistent with fitness effects of deleterious mutations on the order of ~10^−4^ to 10^−5^, 100 to 1000-fold lower than the current *in vitro* estimates [30,31]. While this discrepancy may result from technical difficulties in measuring selection coefficients below 1% with direct *in vitro* competitions, it may also have a biological basis. The mammalian gut is a complex environment that may temporally fluctuate (e.g. variable diet or time of feeding), which could result in temporal variation of the effect of deleterious mutations, with a small time-averaged fitness effect. Indeed, we show in Fig. 2C that the availability of particular resources might buffer the fitness effects of deleterious mutations. Another mechanism for reducing the effects of deleterious mutations is revealed by the nature of the adaptive mutations. As previously observed, adaptation of *E. coli* to the mouse gut involved mutations linked to carbohydrate metabolism, which were under negative frequency-dependent selection, likely created by niche segregation of the different mutants [35,73]. This segregation may reduce competition between lineages and therefore buffer the effect of deleterious mutations. This weak effect of deleterious mutations may be of relevance for a variety of processes both within the gut microbiota (e.g. evolution of antibiotic resistance) and beyond this ecosystem (e.g. cancer evolution). As an example, it has recently been shown that tumour mutational load can be a predictor of survival for some, but not all, types of cancer. Our work suggests that determining the fitness effect of deleterious mutations may be key to understand these differences.

Overall, our work suggests that the evolutionary dynamics of gut commensals may be strongly shaped by beneficial mutations, which increase in frequency but do not fix (partial sweeps), and slightly deleterious mutations that can segregate for long periods of time. The combination of these two effects could allow strains of the same species to coexist, within the gut microbiota, for extended periods with complex temporal dynamics. The few available data on temporal dynamics of genetic polymorphism in species of human gut commensals is starting to confirm this prediction, at least for some species [5–8,59]. As genetic diversity can impact community composition [82], more time series data, similar to that obtained here, is needed towards a full understanding of the selective mechanisms that shape the eco-evolutionary dynamics within the microbiota [83,84].

## Methods

### Bacterial strains

Strains used in this manuscript derive from the commensal *Escherichia coli* K12, substrain MG1655 [85]. These are strains JB19-YFP, JB18-CFP, RR03-YFP and RR04-CFP. JB19-YFP and JB18-CFP were previously described in [35] and have the following genetic background: *galK*::YFP/CFP cm^R^, str^R^ (RpsL^K43R^), *AlaclZYA* and *gatC*::+1bp. The latter mutation was acquired during the first step of adaptation to the mouse gut, as described in [52]. During a second step of adaptation, clones JB18 and JB19 were allowed to adapt to the mouse gut for 24 days. One of the evolved clones (21YFP) acquired two further adaptive mutations: *yjjP/yjjQ*::IS2 and *yjjY/yjtD*::+4bp. To obtain this clone with either CFP or YFP expression, P1 transductions were used to replace the chloramphenicol resistance cassette by an ampicillin resistance cassette [86]. This created clones RR03-YFP and RR04-CFP, with the genetic background *galK*::YFP/CFP amp^R^, str^R^ (RpsL^K43R^), *AlacIZYA, gatC*::+1bp, *yjjP/yjjQ*:: IS2 and *yjjY/yjtD*:: +4bp.

### Mice

All experiments were carried using 6 to 8-week old C57BL/6J male mice raised in specific pathogen-free conditions.

### Ethics statement

All experiments involving animals were approved by the Institutional Ethics Committee at Instituto Gulbenkian de Ciencia (project nr. A009/2010 with approval date 2010/10/15), following the Portuguese legislation (PORT 1005/92), which complies with the European Directive 86/609/EEC of the European Council.

### *In vivo* experimental evolution

To simulate a perturbation of the gut microbiota and allow its invasion by *E. coli*, we provided mice with streptomycin-treated water (5g/L) [52,72] for 24h before gavage with *E. coli* and during one-week post-colonization (Fig. 1A). After this period, mice were given normal water until day 190 post-colonization. On the day of gavage, food and water were removed from the cage for 4 hours. After this period, mice were gavaged with 10^8^ *E. coli* (suspended in 100μl PBS). Four mice were used in this experiment, with two being gavaged with a mixture of RR03-YFP and JB18-CFP and the other two with a mixture of RR04-CFP and JB19-YFP. Clones RR03/RR04 represented 10% of the cells in the inoculum, whereas clones JB18/JB19 represented 90%. These inocula were prepared by growing each clone separately in brain heart infusion medium to OD_600nm_ = 2 (at 37°C on an orbital shaker). After gavage, animals were separated into individual cages and water and food was returned to them. At each sampling point, faecal pellets were collected, weighed and dissolved in PBS, with the remaining volume being stored at −80°C in 15% glycerol. Dilutions were made for plating in LB-agar with streptomycin (100 μg/ml; or streptomycin (100 μg/ml) + ampicillin (100 μg/ml) / streptomycin (100 μg/ml) + chloramphenicol (30 μg/ml)) and plates incubated overnight at 37°C. CFP and YFP colonies were then counted on a fluorescent stereoscope (SteREO Lumar, Carl Zeiss) to estimate total numbers of *E. coli* and the frequency of YFP and CFP clones (Fig. S1).

### Mutator screen and fluctuation tests to estimate mutation rate

#### Mutator screen (Fig. 1B)

Fecal samples isolated from each mouse at day 190 post-colonization were plated as described above. ~90 clones per mouse were isolated and each clone was inoculated into a different well of a 96-well plate with 200μl of liquid LB. Plates were incubated overnight at 37°C on a plate shaker (800 rpm). After the overnight, all clones were frozen at −80°C in 15% glycerol. Stored clones were individually defrosted into 200μl LB in a 96-well plate and incubated overnight as described above. 10μl of each culture were removed to quantify the total number of bacteria by FACS, which was generally >10^8^ per well (only 10/430 samples were between 10^7^ and 10^8^). The remaining volume was plated in LB-agar plates with nalidixic acid (40 μg/ml) to determine the number of resistant mutants that emerged during growth. Plates were incubated as described above.

#### Fluctuation tests to estimate mutation rate (Fig. 1C and 2B)

Individual clones, stored at −80°C, were defrosted into PBS, plated and grown overnight at 37°C. We then picked individual colonies to PBS and used FACS to adjust the inoculum size to 2000 cells per replicate. Cells were then inoculated into 200μl of liquid LB and grown in a 96-well plate for 20-24h (incubated at 37°C, 800 rpm in a plate shaker). For some clones this was done in 1000μl of liquid LB with growth in 96-deep well plates. At this time point, 10μl of each culture were removed and used to make dilutions to quantify the total number of cells by plating in LB-agar plates. The remaining culture volume (and/or a 10^−1^ dilution) was then plated in LB plates with rifampicin (100 μg/ml) or nalidixic acid (40 μg/ml) to quantify the number of resistant mutants that emerged during growth. All fluctuation tests were repeated in two independent blocks. The total number of colonies and the number of resistant colonies were then used to estimate mutation rate (and 95% confidence intervals) with FALCOR (using the Ma-Sandri-Sarkar maximum likelihood estimator; http://www.keshavsingh.org/protocols/FALCOR.html) [87–91]. Significant differences in mutation rates between clones were identified from non-overlapping 95% confidence intervals.

### Whole-genome sequencing and analysis

#### Sequencing

19 clones were sequenced from mouse 1 (where mutators emerged; 18 CFP and 1 YFP), 3 from mouse 2, 3 from mouse 3 and 4 from mouse 4. Bacterial clones were defrosted into liquid LB and grown overnight or into PBS, plated in 2-3 LB agar plates and incubated for 24h at 37°C. After pelleting the cells, we extracted DNA following the procedure described in [92]. DNA library construction and sequencing were performed by the IGC genomics facility. Each clone was paired-end sequenced on an Illumina MiSeq Benchtop Sequencer. Standard procedures produced data sets of Illumina paired-end 250 bp read pairs, with mean coverage of 32-fold (19 to 45-fold per clone). The *fastq* files for all sequenced clones will be deposited at the NCBI SRA.

#### Variant calling

*breseq* v0.26 [93], with default parameters, was used to map reads and identify mutations, using the *E. coli* str. K-12 substrain MG1655 genome as reference (NCBI accession: NC_000913.2). Mutations were then determined by comparison with the ancestrals (JB18, JB19, RR03, RR04, sequenced for [35]). The reads that *breseq* failed to match to the reference genome were assembled into contigs with *SPAdes* v.3.13 [94]. All contigs with coverage >3 were then manually annotated via BLAST [95]. All annotated contigs matched the cassettes carrying the antibiotic resistance plus the fluorescent protein (with a single exception, a likely contaminant), suggesting that horizontal gene transfer did not occur during this experiment.

#### Phylogeny

The phylogenetic tree in Fig. 3B and S10 is a minimal evolution tree [96], constructed with the R package *ape* (v5.2) [97] by using a matrix of the raw mutational distance between clones and plotted with the *ggtree* package (v1.14) [98,99].

#### Parallel mutational targets across non-mutator clones (for Fig. 4B)

These were identified by finding the mutational targets (genes and operons) that independently acquired mutations in more than one non-mutator clone (Fig. S11). As mutations are random and only 3 to 8 mutations accumulated in non-mutator clones, it is very unlikely that we would find the same gene/operon being independently mutated if such mutations did not carry a fitness benefit [100]. Using this strategy, we identified 13 mutational targets that are candidates for adaptation, 12 of which were mutated in clones isolated from different mice and a single one that acquired independent mutations in two clones isolated from the same animal.

#### Parallel mutations across mutator clones (within mouse 1; for Fig. 4B)

As mutator clones have long terminal branches (Fig. 3B), most of the evolutionary history of each clone is independent of the others. Thus, the mutations accumulating at these terminal branches can be used to identify parallel mutational targets. However, as the mutation rate is up to 1000-fold higher than non-mutator clones, there is the possibility that some mutational targets would accumulate multiple independent mutations just by chance (e.g. longer genes have a higher per gene probability to acquire a mutation). Thus, here we used a statistical approach to identify which genes accumulated more SNPs (at the terminal branches) across independent clones than would be expected by chance. We focused on SNPs because these are the large majority of the observed mutations (Fig. S10) and on genes because these are straightforward to define at both the sequence and functional levels. The statistical approach we used is similar to the one in [80]. Briefly, we first simulated 100 datasets in which we randomly distributed the 3245 independent SNPs (synonymous, non-synonymous and intergenic) that we identified in the mutators. This randomization process was done taking into account the observed number of each specific substitution (*i.e*. if it is A →T or G →C, etc). From these simulations, we kept the non-synonymous mutations and calculated a G score for each gene in each simulation. This involved first calculating the expected number of mutations (*E_i_*), as:

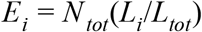

where *N_tot_* represents the total number of non-synonymous SNPs (per simulation), *L_i_* is the length of gene *i* and *L_tot_* is the total size of the coding genome. The G score for gene *i* can then be calculated as:

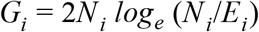

where *N_i_* is the number of non-synonymous SNPs at gene *i*. When *N_i_*=0, we set *G_i_*=0. Using the same approach, we calculated *G* scores for the observed mutations at each of the mutated genes. We could then calculate a *Z* score for each gene *i* at which mutations were observed: 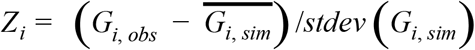, where *G_i, obs_* and *G_i, sim_* are the G score for the observed mutations and the simulations at gene *i*, respectively (overbar and *stdev* indicate mean and standard deviation). The *Z_i_* scores were then used to calculate Benjamini-Hochberg corrected *p*-values. Furthermore, we applied two additional conservative criteria: 1) we divided the observed *N_i_* and *N_tot_* by 2 before estimating *G_i, obs_*. This takes into account that mutation rates can vary along the genome, in general by 2-fold [101–103]. Dividing *N_i_* and *N_i, tot_* by 2 aims to control for the possibility that significant genes are in regions where mutation rate is highest. 2) We included only genes that were mutated in at least 3 independent clones, as this leads to a similar probability of a mutation being parallel as for non-mutator clones, where two independent mutations are considered (*i.e*. for non-mutator: *U*^2^ = (5×10^−10^)^2^=2.5×10^−19^; for mutator: *U*^3^ = (5×10^−7^)^3^=1.25×10^−19^). This analysis led to the identification of the parallel mutational targets described in Table S3.

#### dN/dS estimates

As above, we used the mutations at the terminal branches to estimate dN/dS for each mutator clone, by taking the ratio of the non-synonymous to synonymous SNPs. Given that the probability of a SNP to be non-synonymous is not uniform across the different types of substitutions, we also estimated the expected dN/dS, given the observed mutational spectra of each clone (using a similar approach to the described in [104]). dN/dS values, shown in Fig. S10, are normalized by the expected value and *p*-values are obtained with a binomial test.

#### DNA polymerase Ill structure (Fig. 2A)

obtained from rcsb.org (PDB ID: 5M1S) [40,105] and edited with PyMOL to display mutations [106].

### Allelic reconstructions

The mutations *DnaQ*^L145P^ (C<u>T</u>C→C<u>C</u>C) and DnaE^T771S^ (<u>A</u>CG→<u>T</u>CG) were constructed by pORTMAGE recombineering [107] in the ancestral (RR04-CFP) background, with the pORTMAGE-3 plasmid (carrying the kanamycin resistance cassette; oligomers are listed in Table S5). The double mutant was constructed with the same method, by inserting the DnaQ^L145P^ mutation in the *dnaE* background. The presence of the desired mutations was confirmed by PCR and Sanger sequencing, after which we grew each clone in LB to lose the pORTMAGE plasmid (this was confirmed by streaking the clones in LB-agar plates with or without kanamycin, 100 μg/ml).

### *In vivo* and *in vitro* mutation frequency temporal dynamics

To measure the *in vivo* mutation frequency (Fig. 2D-E), we inoculated different mice with one of three different clones: Ancestral (RR04), a DnaQ^L145P^ single mutant (*dnaQ*; randomly chosen from the clones with highest mutation rate) and a DnaQ^L145P^ + DnaE^T771S^ double mutant (*dnaQ*; chosen as for single *dnaQ* clone). All clones carry the CFP marker. Mice were provided with streptomycin-treated water (5 g/L) for 7 days before gavage with *E. coli*. 4 hours before gavage, food and water were removed from the cages. After gavage, animals were separated into individual cages and food and normal water (without streptomycin) was returned to them. To prepare the inoculum for these experiments, we defrosted each clone into PBS and plated in LB agar plates. After overnight incubation at 37°C, colonies were scrapped into PBS and OD_600nm_ adjusted to 2. 100 μl of this suspension was then used to inoculate each mouse (*i.e*. 10^8^ *E. coli*). Independent replicate inocula were used to colonize independent mice. This procedure for preparing the inocula is different from the above because we wanted to minimize the strength of selection for compensatory mutations, which might reduce the mutation rate and/or the growth effects of the mutations (growth in liquid media, where competition is global, is known to lead to more rapid adaptation than in solid media, where competition is local; [108]). Faecal samples were then obtained at 6h and every 24h after gavage for 4 days and suspended in PBS. These were directly plated in LB-agar plates with streptomycin and rifampicin to quantify the number of *de novo* rifampicin resistant mutants and dilutions were made to plate in LB-agar plates with streptomycin to quantify the total number of *E. coli* (all clones are streptomycin-resistant). These two numbers were then used to estimate the equilibrium mutation frequency, which is proportional to mutation rate (as described in the results section).

As we had estimated mutation rate *in vitro* with fluctuation tests, we carried a control experiment to understand if the equilibrium mutation frequency was similar between the *in vitro* and *in vivo* conditions. For this, we prepared the inocula of the three different clones as described above, but started cultures with 10^6^ *E. coli* in 3 ml liquid LB (in 15ml tubes). Cultures were incubated for 24h at 37°C in an orbital shaker. Every 24h, cultures were diluted 1000-fold and allowed to grow until saturation (population size ~ 3 to 5×10^9^; 10 generations per day). This procedure was repeated for 4 passages. Every day, the cells were plated as described for the *in vivo* experiment. The results of this experiment showed that the mutation frequency for all mutants was similar between the *in vitro* and the *in vivo* conditions (Fig. S5).

We used linear mixed models to analyze temporal of mutation frequency and bacterial density, with replicate as a random effect. If needed, data were transformed to meet assumptions made by parametric statistics.

### *In vitro* growth assays in different media

Bacteria were defrosted into 150μl of liquid LB, grown overnight in a 96-well plate and incubated at 37°C in a plate shaker (at 800 rpm). OD_600nm_ was then adjusted to 0.01, cultures diluted 1:100 and 5μl spotted in agar plates and incubated at 37°C for 48h. After incubation, growth of each clone was imaged on a fluorescent stereoscope (SteREO Lumar, Carl Zeiss). The same inocula were tested across all media. The media used were LB and M9 minimal media with: 0.4% glucose, 0.4% sorbitol and 0.4% glucose plus the 20 proteinogenic amino acids (each at 0.05 mM) (Fig. 2C).

### Statistical analysis

All analysis were performed in R, version 3.5.1 [109], using the statistical methods described in the above sections.

### Simulations of Muller’s ratchet

All simulations were run in Mathematica [110]. The annotated simulation code will be made available in an online depository.

Simulations of Muller’s ratchet with fixed effects of deleterious mutations were implemented similar to those in [49]. Beginning with a mutation-free population of 10^6^ individuals, each generation mutations were drawn from a Poisson distribution with parameter *μ* (mutation rate). The fitness of each individual was computed as *w_i_*=(1-*s_d_*)^*i*^, where *i* is the number of mutations carried, and *s_d_*>0 is the deleterious selection coefficient. The next generation of the population was obtained by sampling 10^6^ individuals from the current population according to probabilities weighed by the individuals’ absolute fitness. After 1000 and 2000 generations, respectively, we recorded the minimum number of mutations accumulated within the population (*i.e*., number of mutations in the least-loaded class), the mean and standard deviation of the number of mutations carried across the whole population, and the mean and standard deviation of fitness within the population.

For simulations with a continuous distribution of fitness effects (Fig. S8), upon mutation, an individual fitness effect per mutation was drawn from an exponential distribution with parameter 1/*s_d_*. To consider beneficial mutations, a fraction *f* of these mutations were randomly chosen and their selection coefficient multiplied with −1, rendering them beneficial.

For a given combination of *U_d_* and *s_d_*, we considered that the number of mutations in the simulations fitted those in the experiments if the number of observed mutations for any clone was contained within the mean±standard deviation of any simulation (Fig. 3A and Fig. S6–8).

For simulations of two subpopulations with different mutation rates (Fig. S9), a marker locus was implemented that determines the mutation rate within the subpopulation built by its carriers. The number of individuals of subpopulation 1 in the next generation *N*_1_ (*t* + 1) was sampled from a binomial distribution with parameters N=10^6^ and 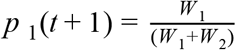, where *W_k_* is the total fitness of subpopulation *k* at generation *t* (*i.e*. the sum of all current individual fitnesses in this subpopulation). The number of individuals in subpopulation 2 was then determined as *N*_2_(*t* + 1) = *N* – *N*_1_(*t* + 1).

Finally, for simulations with negative frequency-dependent selection (Fig. 4D), the number of individuals of subpopulation 1 in the next generation *N*_1_(*t* + 1) was sampled from a binomial distribution with parameters N=10^6^ and 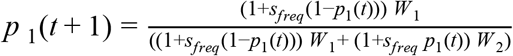, where *S_freq_*=0.1.

## Acknowledgements

We thank Daniela Gülereşi for experimental assistance and Brian Charlesworth for highlighting to us how weak the deleterious effects of mutations may be. We thank Ana Sousa, Roberto Balbontín and Ana-Hermina Ghenu for helpful comments on the manuscript. We acknowledge the technical support of the following IGC facilities: Genomics Unit, Rodent facility, Flow Cytometry facility and Advanced Imaging facility.

---

**Figure S1.**
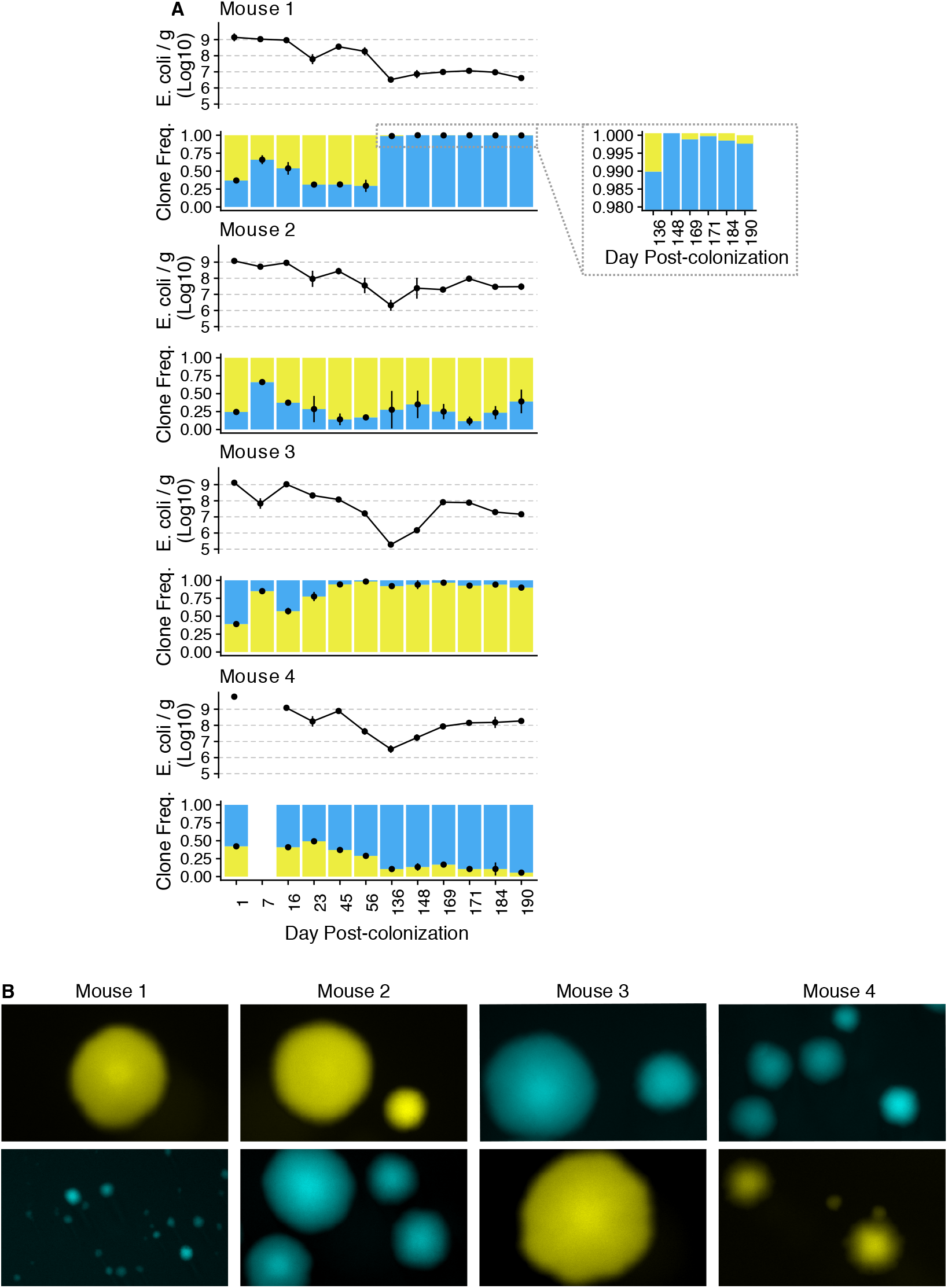
Temporal dynamics of *E. coli* densities and frequency of two clones over 190 days (A) and colony size variation at day 190 (B). Mouse 1 is where mutators emerged. Clones used to colonize mice 1 and 2: YFP, *gatC*::+*C* (yellow); CFP, *gatC*::+C, *yjjP/yjjQ*::IS2, *yjjY/yjtD*::+TTAT (blue). Clones used to colonize mice 3 and 4 have the same genotype, but the markers are swapped. Specifically, mice3 and 4 were colonized with: CFP, *gatC*::+C (blue); YFP, *gatC*::+C, *yjjP/yjjQ*::IS2, *yjjY/yjtD*::+TTAT (yellow).

**Figure S2.**
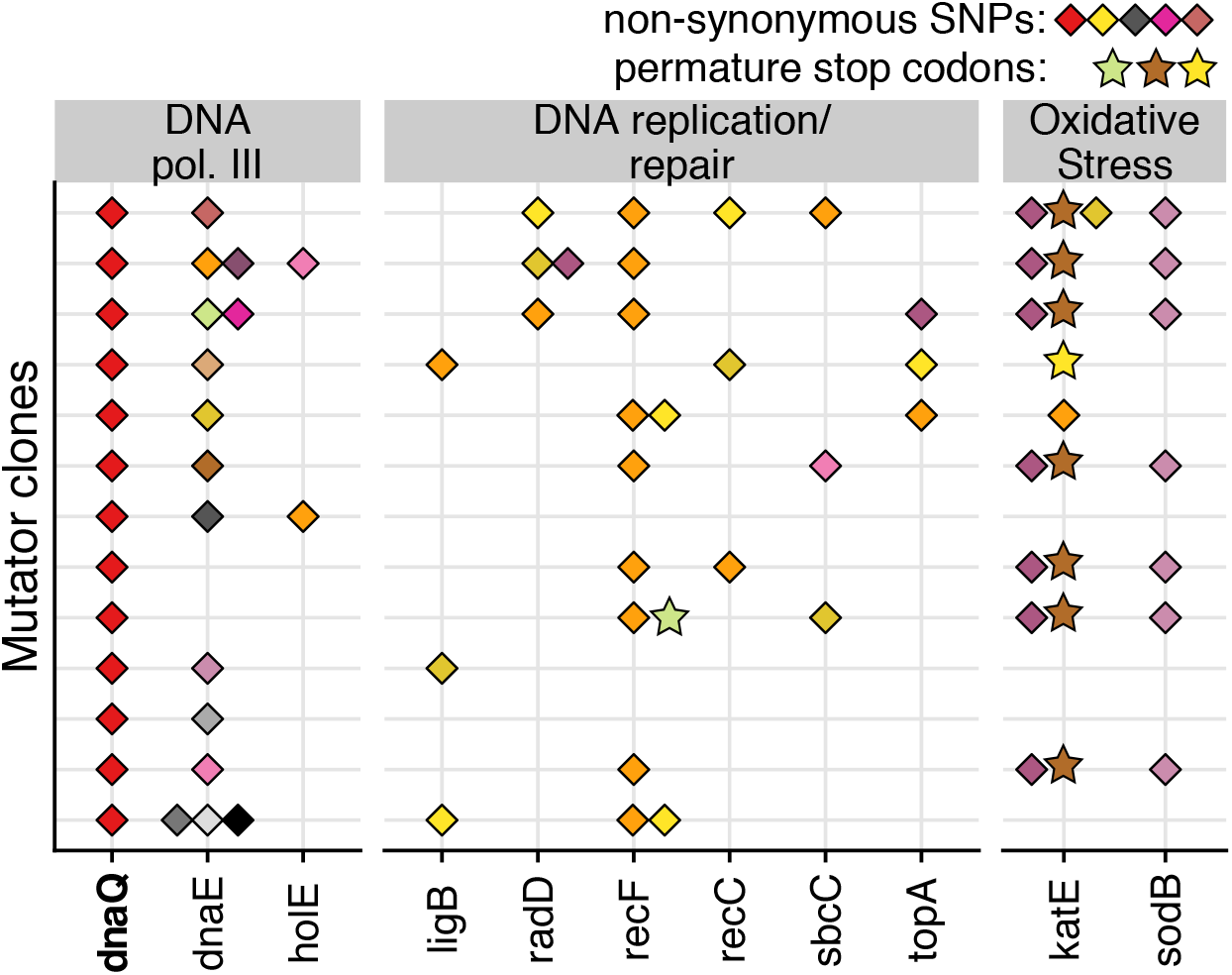
Mutators accumulate multiple mutations in genes that can affect mutation rate. Each row represents a clone and points with different colors represent different alleles. Genes were identified from Ecocyc (by looking at genes involved in DNA repair and replication) [111] and from [41, 112–115].

**Figure S3.**
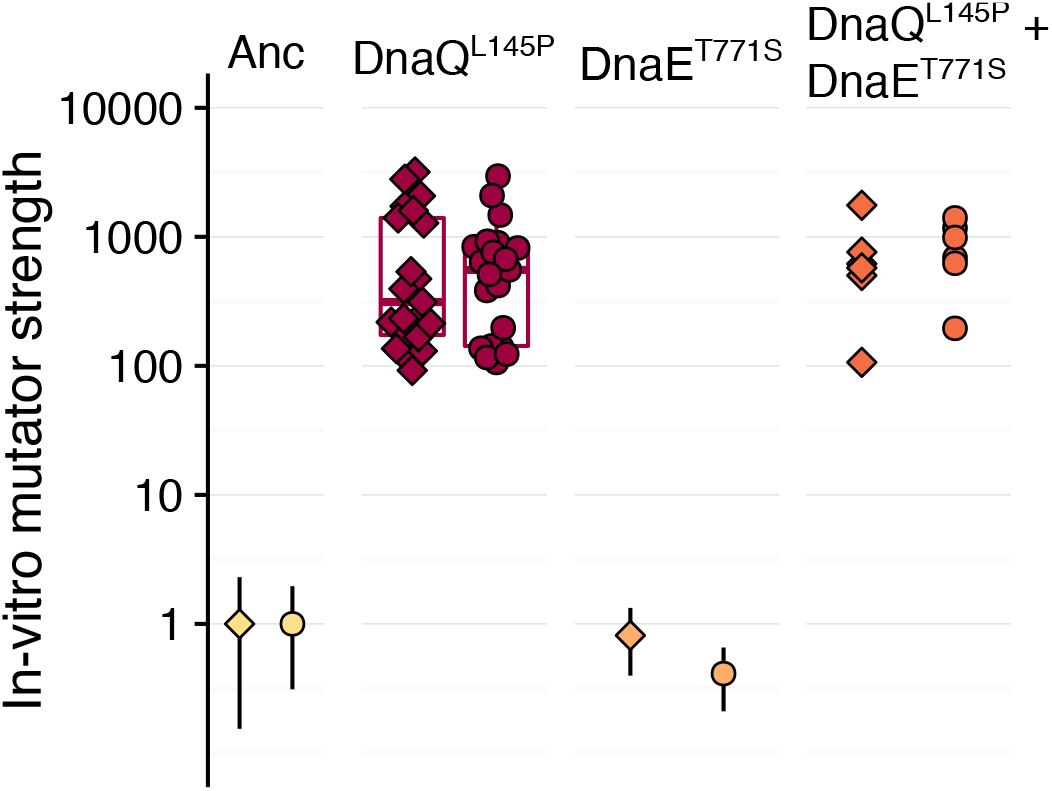
*In vitro* mutator strengths, estimated from fluctuation tests, for the Anc, DnaQ^L145P^, DnaE^T771S^, DnaQ^L145P^+DnaE^T771S^. Data for Anc, DnaQ^L145P^ and DnaQ^L145P^+DnaE^T771S^ are the same as in Fig. 2B. These are repeated here to enable visual comparison. For ease of visualization, 95% confidence intervals are only shown for the ancestral and DnaE^T771S^, but none of the 95% CI of the DnaQ^L145P^ or the DnaQ^L145P^+DnaE^T771S^ mutants overlap with either the ancestral or the DnaE^T771S^ mutant (See Table S1 for mutation rates and 95% CI).

**Figure S4.**
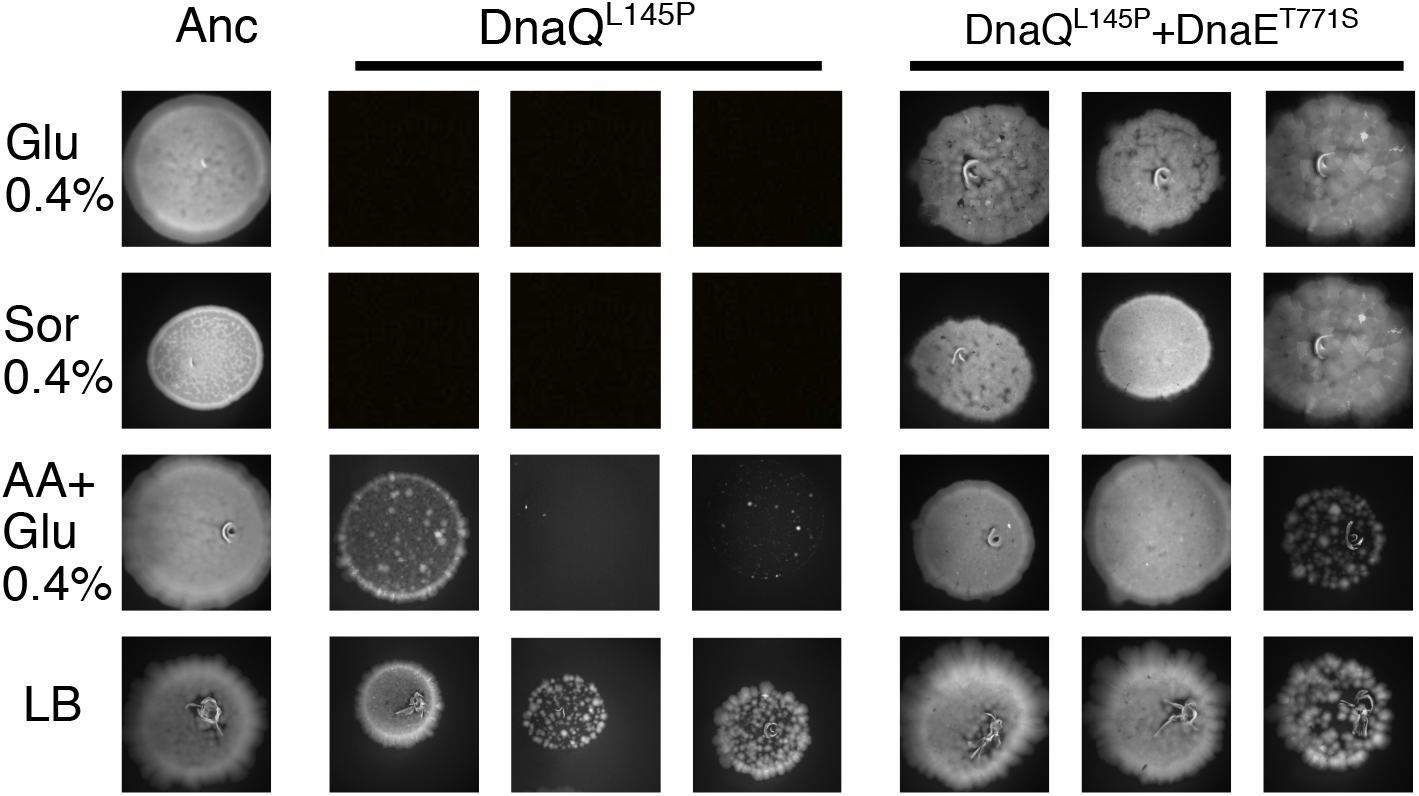
*In vitro* growth capacity of additional independent clones of the mutants DnaQ^L145P^ and DnaQ^L145P^+DnaE^T771S^. Spot assay in M9 minimal media with 0.4% glucose (Glu 0.4%), 0.4% sorbitol (Sor 0.4%); 0.4% glucose plus the 20 proteinogenic amino acids (each at 0.05 mM; AA+Glu 0.4%) or LB.

**Figure S5.**
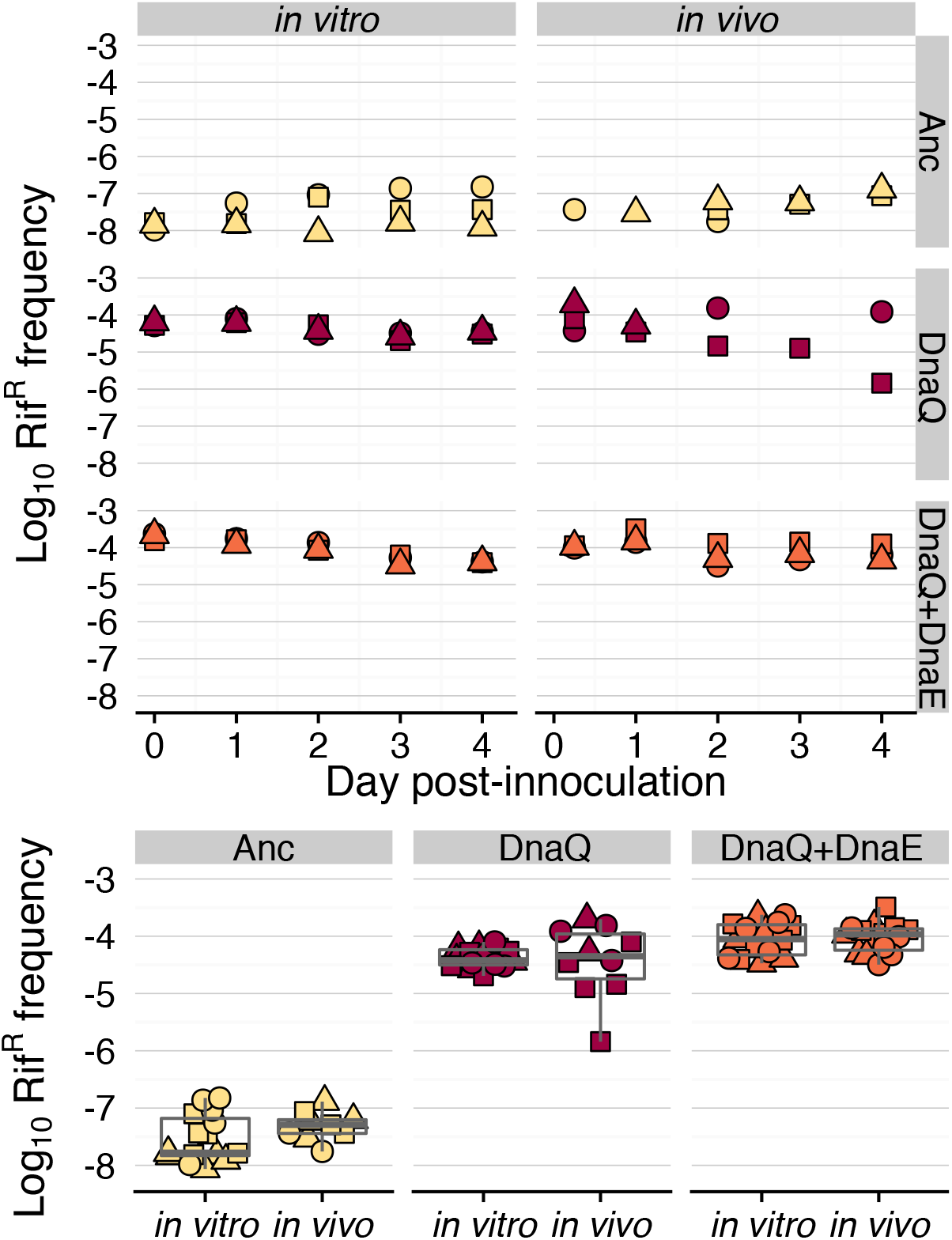
Mutation frequency has similar dynamics in vitro and in vivo. Top: Temporal dynamics for the frequency of rifampicin-resistant mutants during an in vitro propagation in LB (left) and *in vivo* colonization of the mouse gut (right) with the Ancestral, DnaQ^L145P^ mutant and DnaQ^L145P^+DnaE^T771S^ double mutant. Bottom: Boxplots showing the pooled data points across all time points (colors represent different clones and shapes represent different replicates). Using a linear mixed model (with replicate as random effect), we find that there is no significant effect of either the interaction between clone and experiment (**χ**^2^_2_=0.60, p=0.74) or of experiment (**χ**^2^_1_=0.29, p=0.59), with the clone being the only significant effect (**χ**^2^_1_=42.06, p<0.0001).

**Figure S6.**
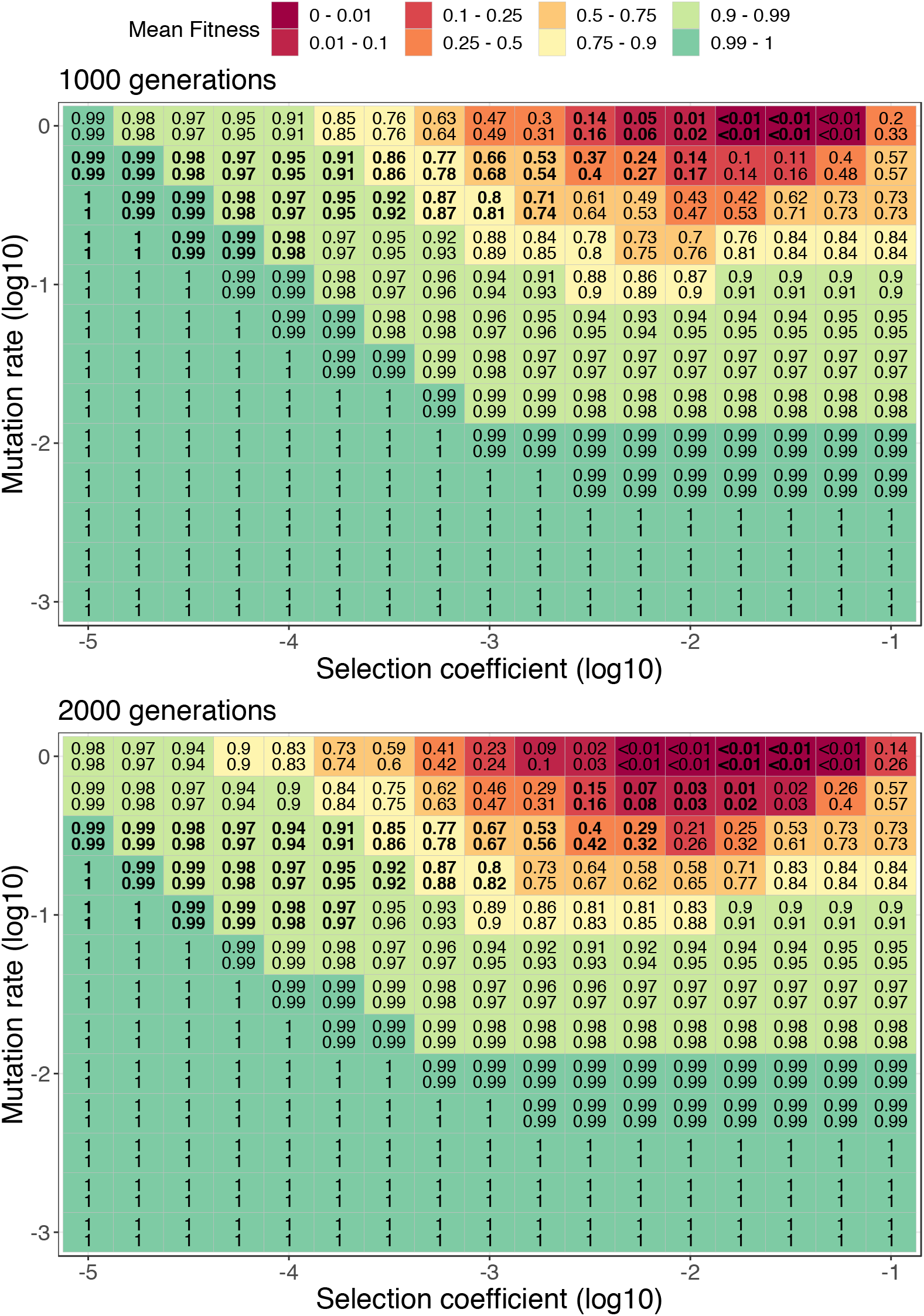
Mean fitness in Muller’s ratchet simulations with deleterious mutations of fixed effect and a clonal population size of 10^6^ (n=10 simulations). Colour gradient indicates the mean absolute fitness at 1000 and 2000 generations. For each simulation, the mean fitness of the population, at 1000 and 2000 generations, was calculated. The numbers inside each square (i.e. selection coefficient/mutation rate combination) indicate the minimum (top) and maximum (bottom) mean fitness that was observed across the 10 simulations.

**Figure S7.**
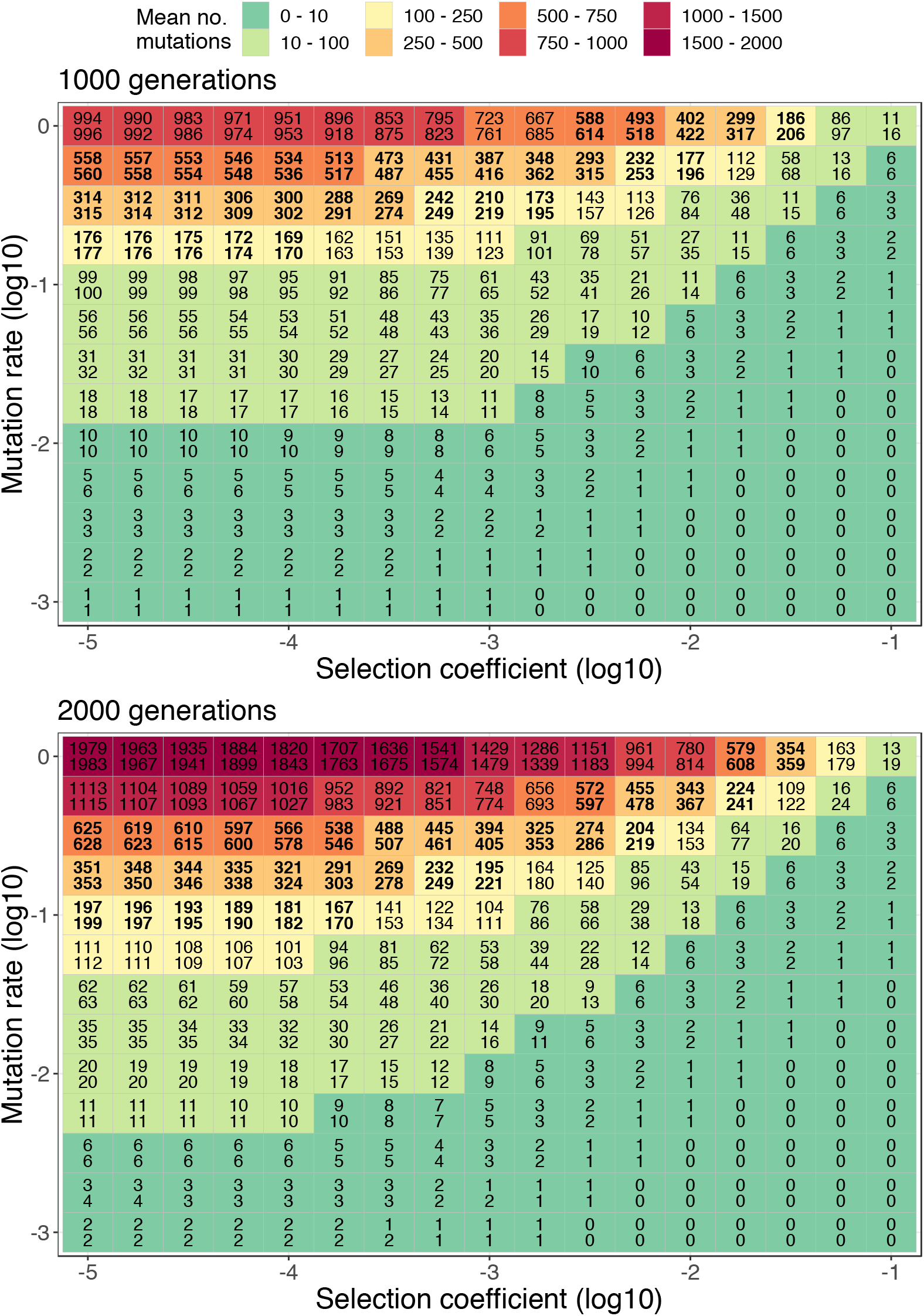
Mean number of mutations in Muller’s ratchet simulations with deleterious mutations of fixed effect and a clonal population size of 10^6^ (n=10 simulations). Colour gradient indicates the mean number of mutations at 1000 and 2000 generations. For each simulation, the mean number of mutations of the population, at 1000 and 2000 generations, was calculated. The numbers inside each square (i.e. selection coefficient/mutation rate combination) indicate the minimum (top) and maximum (bottom) mean number of mutations that was observed across the 10 simulations.

**Figure S8.**
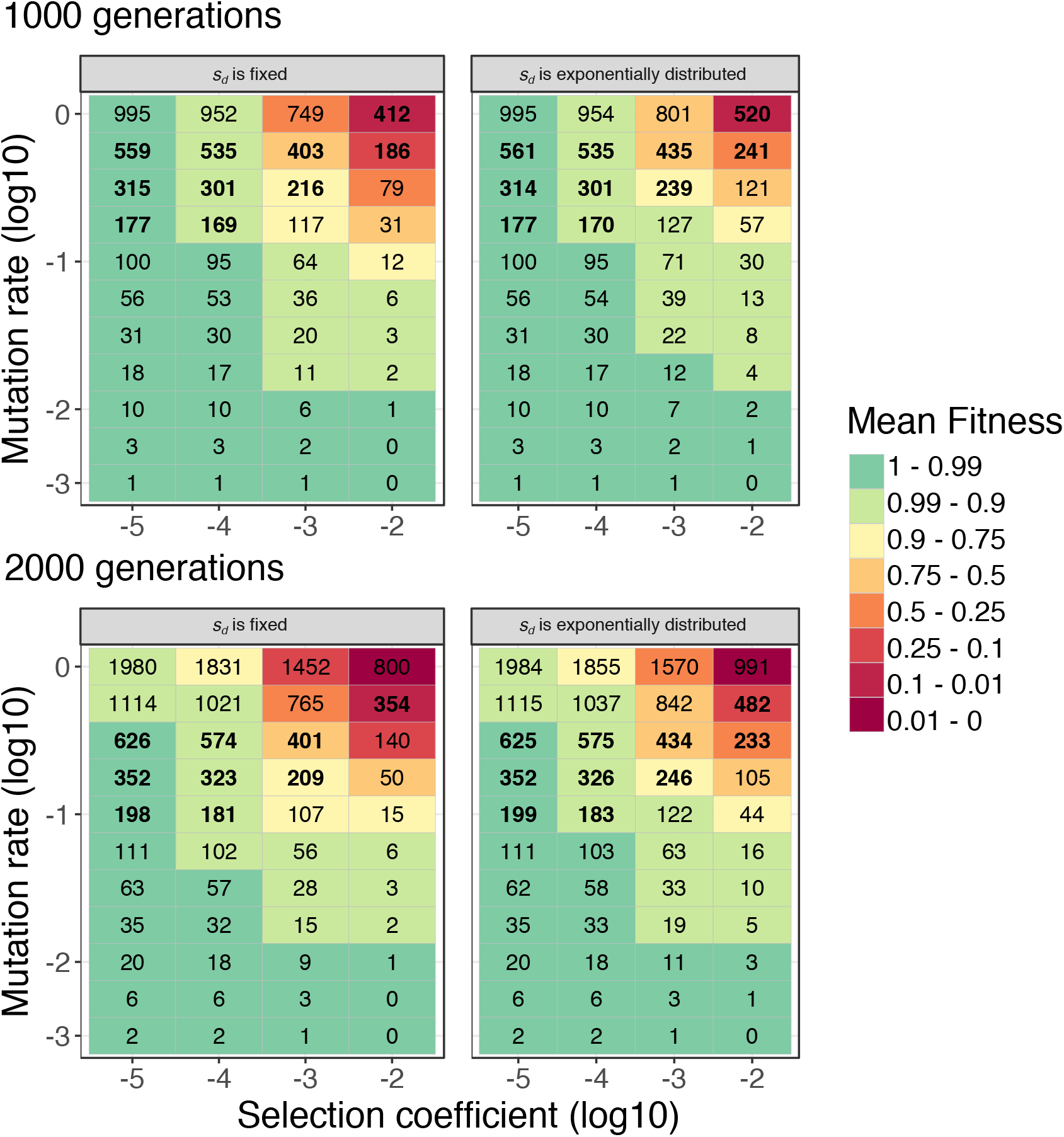
Weak effects of deleterious mutations (*s_d_*) are necessary for strong mutators to escape Muller’s ratchet, independently of whether *s_d_* is fixed or exponentially distributed. Mean fitness and mean number of mutations in Muller’s ratchet simulations with deleterious mutations of fixed (left) or exponentially distributed effect (right). Colour gradient indicates the mean absolute fitness after 1000 (top) and 2000 generations (bottom) for different combinations of *s_d_* and mutation rate (*U_d_*). Numbers in bold indicate the parameter space for which there is a fit between the observed (Fig. 2B) and the simulated number of mutations. The observed variation in the number of mutations is only possible if *s_d_*<10-4 and if there is mutation rate variation. Population size = 10^6^. Fixed *s_d_*: averages of n=10 simulations; Exponentially distributed *s_d_*: n=1 simulation.

**Figure S9.**
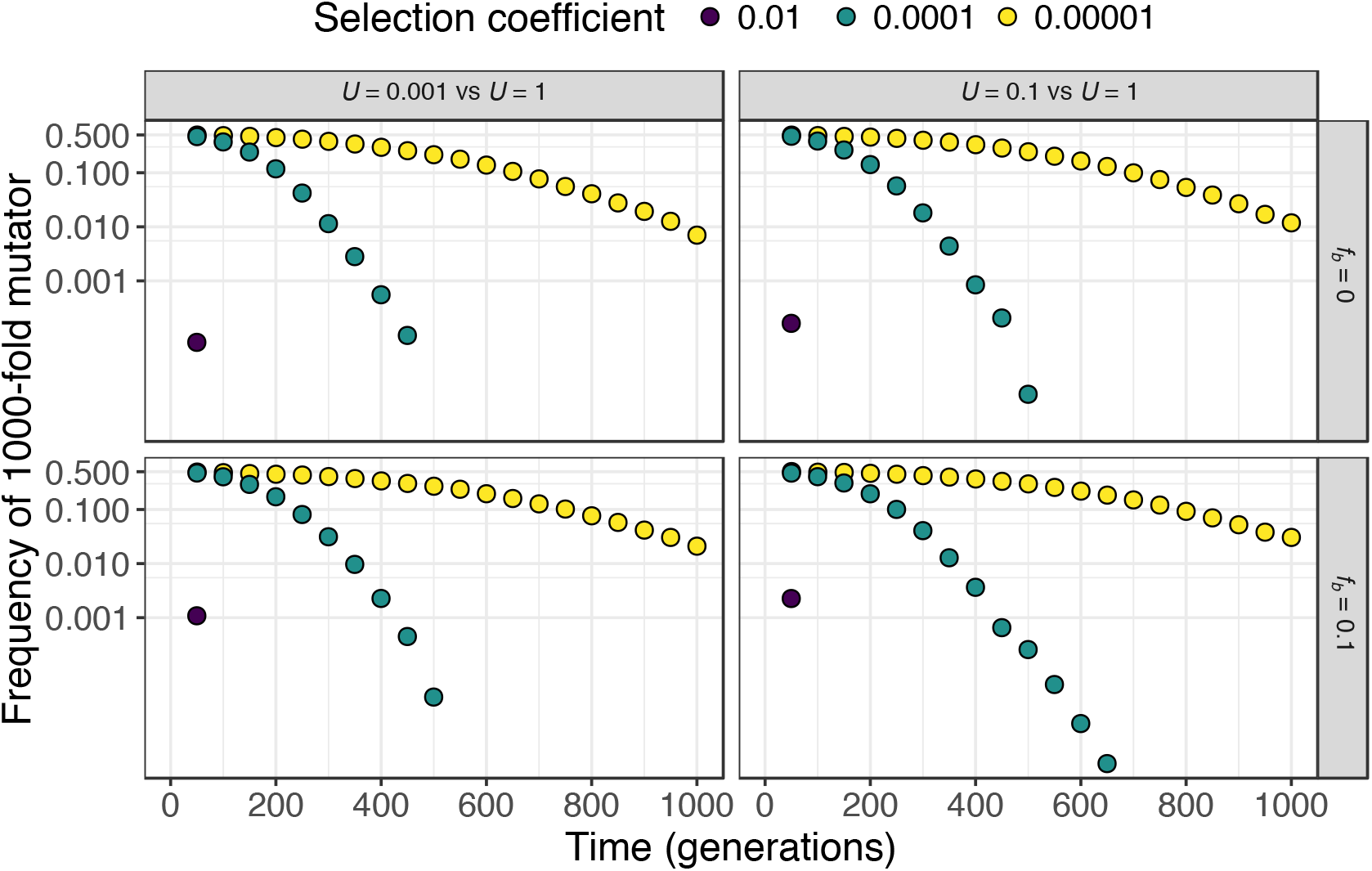
Weak effects of deleterious mutations allow 1000-fold mutators to coexist with clones with lower mutation rates and this is largely independent of the presence of beneficial mutations. Frequency of a clone with *U* = 1 (1000-fold mutator), when in competition with a clone with *U* = 0.001 (wild-type mutation rate; left) or a clone with *U* = 0.1 (100-fold mutator; right), either in the absence of beneficial mutations (*f_b_* = 0; top) or with a 10% fraction of beneficial mutations (*f_b_* = 0.1; bottom) and with different selection coefficients (population size of 10^6^; n=1 simulation per parameter combination). Simulations involve competing pairs of clones with different mutation rates, each starting at 0.5 frequency. Beneficial and deleterious mutations are drawn from a symmetrical exponential distribution and thus the mean fitness effect of beneficial and deleterious mutations is the same, with only the fraction of beneficials changing. Mutators coexist for 50, 450-650 or 1000 generations for *s_d_* values of 0.01, 0.0001 or 0.00001, respectively.

**Figure S10.**
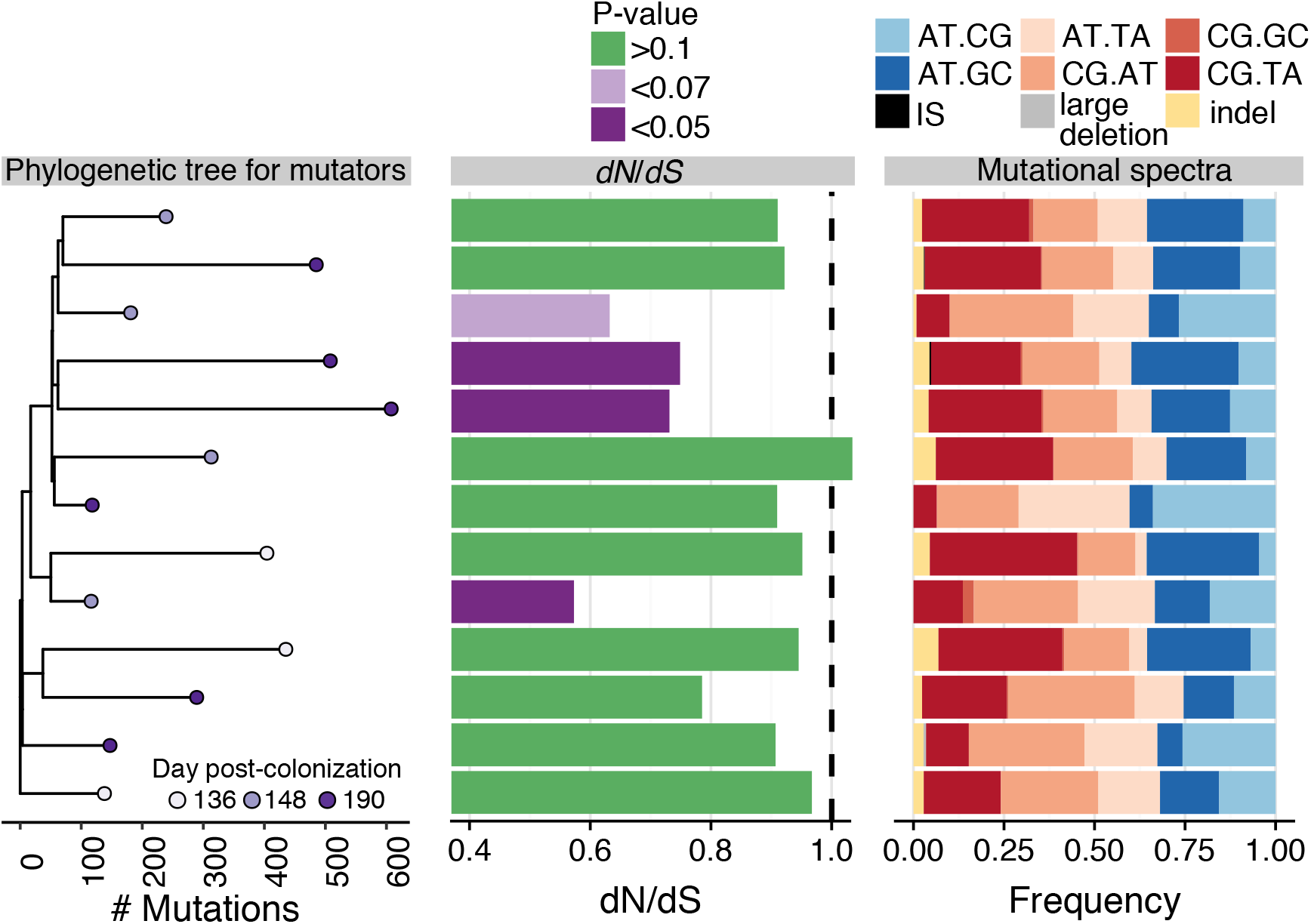
Molecular phenotypes of mutator clones. A) Phylogenetic tree for the mutator clones sequenced from mouse 1. B) dN/dS with colours highlighting whether this is significantly different from neutral expectation, C) mutational spectra. Mutations accumulated at the tips were chosen for this analysis as a way to avoid counting the same mutation multiple times.

**Figure S11.**
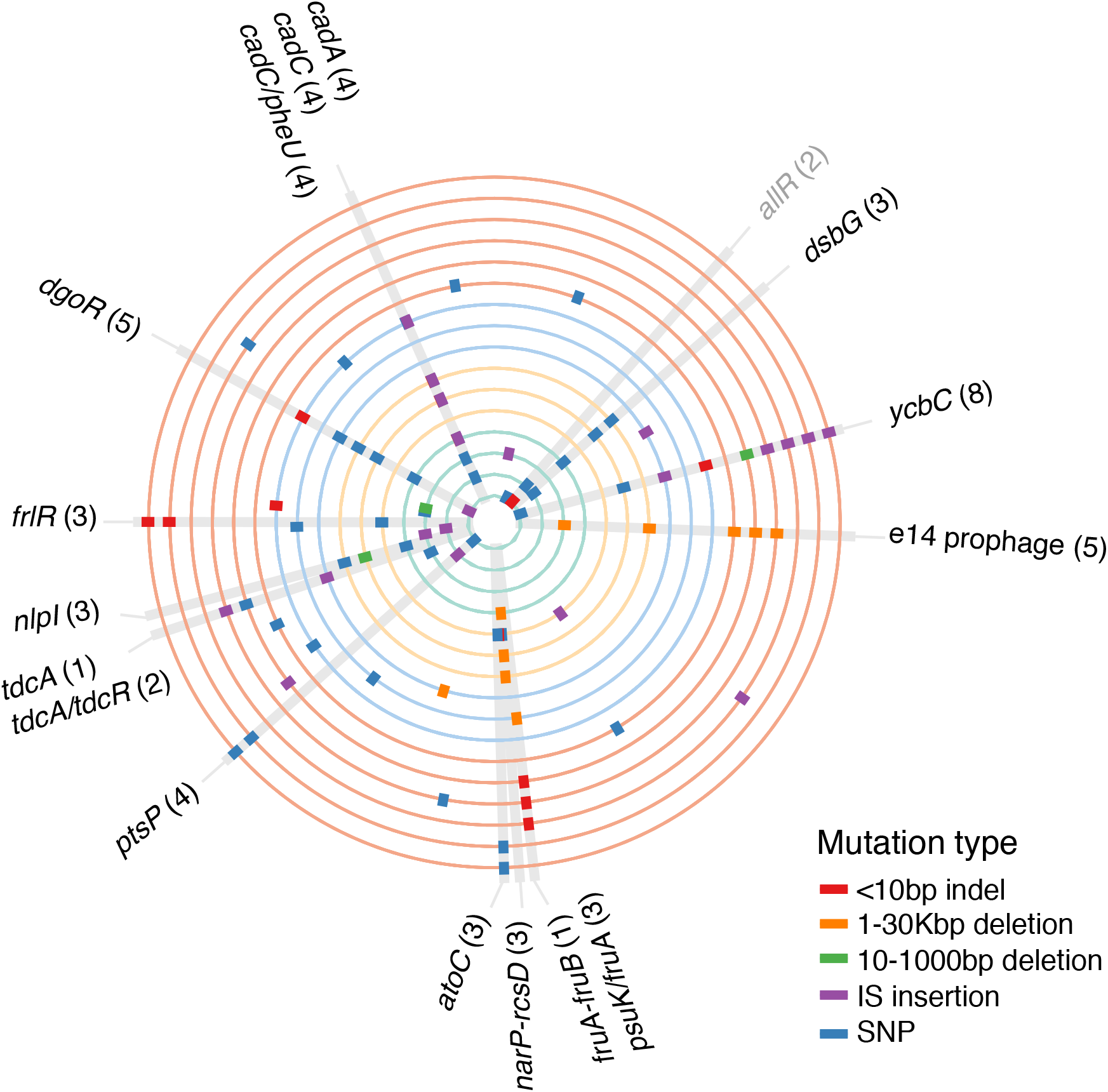
Mutations observed across non-mutator clones isolated from the four different mice (3 to 6 clones per mouse). Circles represent genomes, different coloured circles represent different mice (orange - mouse 1, blue - mouse 2, yellow - mouse 3, green - mouse 4). Grey rectangles highlight mutations that were parallel across different mice (gene labels in black) or for which multiple alleles were found within the same mouse (gene labels in grey). Small rectangles crossing a particular circle indicate mutations, with colours representing different mutation types. Note that for the figures in the main text, we always show 5 nonmutator clones from mouse 1 and there are 6 in this figure. This is because we sequenced one clone at day 190 from the YFP background. As this is not genetically related to the ancestral from which the mutators emerged, we do not show this clone in the main figures.

